# Deep Gaussian Process with Uncertainty Estimation for Microsatellite Instability and Immunotherapy Response Prediction Based on Histology

**DOI:** 10.1101/2024.11.01.621561

**Authors:** Sunho Park, Morgan F. Pettigrew, Yoon Jin Cha, In-Ho Kim, Minji Kim, Imon Banerjee, Isabel Barnfather, Jean R. Clemenceau, Inyeop Jang, Hyunki Kim, Younghoon Kim, Rish K Pai, Jeong Hwan Park, Jewel J. Samadder, Kyo Young Song, Ji-Youn Sung, Jae-Ho Cheong, Jeonghyun Kang, Sung Hak Lee, Sam C. Wang, Tae Hyun Hwang

## Abstract

Determining tumor microsatellite status has significant clinical value because tumors that are microsatellite instability-high (MSI-H) or mismatch repair deficient (dMMR) respond well to immune check-point inhibitors (ICIs) and oftentimes not to chemotherapeutics. We propose MSI-SEER, a deep Gaussian process-based Bayesian model that analyzes H&E whole-slide images in weakly-supervised-learning to predict microsatellite status in gastric and colorectal cancers. We performed extensive validation using multiple large datasets comprised of patients from diverse racial backgrounds. MSI-SEER achieved state-of-the-art performance with MSI prediction, which was by integrating uncertainty prediction. We achieved high accuracy for predicting ICI responsiveness by combining tumor MSI status with stroma-to-tumor ratio. Finally, MSI-SEER’s tile-level predictions revealed novel insights into the role of spatial distribution of MSI-H regions in the tumor microenvironment and ICI response.

## 1 Introduction

Patients whose cancers are microsatellite instability-high (MSI-H)/deficient in mismatch repair (dMMR) have better outcomes than patients with microsatellite stable (MSS) tumors [1, 2]. Additionally, MSI-H/dMMR tumors are highly sensitive to immune checkpoint inhibitors (ICIs) and may not respond to traditional chemotherapy [3, 4, 5].

MSI testing is recommended for all newly diagnosed gastric [6] and colorectal cancers [7, 8]. Current methods of testing include immunohistochemistry (IHC) for detecting MMR status and polymerase chain reaction (PCR) for determining MSI status. However, these assays are time-intensive and costly, and many patients do not undergo the recommended molecular profiling [9]. Numerous studies have demonstrated the feasibility of using deep learning algorithms to analyze hematoxylin and eosin (H&E)-stained whole-slide images (WSIs) to predict MSI status [10, 11, 12, 13, 14, 15, 16, 17, 18, 19, 20, 21, 22].

Thus, incorporating artificial intelligence (AI) into the clinical workflow may provide cost-efficient and widely accessible MSI testing. The adoption of AI-based MSI-status prediction into routine clinical practice requires extensive validation in large, diverse patient cohorts. The inclusion of heterogeneous patient cohorts is particularly important as there may be biological differences associated with race and ethnicity [23, 24, 25]. A recent study showed that a model trained on a predominantly non-Hispanic White patient cohort with gastric cancer, performed poorly when it was tested on samples from Asian patients [10]. These data highlight the fundamental need to validate novel clinical tools across diverse populations.

The capability to quantify uncertainty in predictions is not only crucial to enhance a model’s predictive accuracy, but it also may guide physicians to make more informed decisions. Cases with high predictive uncertainty will require nuanced decision-making by human experts. While numerous deep learning methods, including convolutional neural networks (CNNs) and vision-transformer-based methods, have been applied to MSI status prediction problems, most do not capture the uncertainty in the prediction as point estimation methods. There is a need for a prediction model to not only deliver accurate predictions, but also has the capability to quantify the uncertainty of the predictions. Finally, previously reported algorithms also focus solely on MSI prediction without providing insights into ICI responsiveness, which limits their clinical utility.

To address these challenges, we propose MSI-SEER, which utilizes deep Gaussian processes (DGPs) [26] in weakly supervised learning, a form of inexact supervision tasks [27], to analyze H&E-stained WSIs. MSI-SEER predicts MSI-H status by first calculating the probability of being MSI-H for each tile within a WSI, then aggregating these tile-level probabilities to assess the overall MSI-H status of the slide. This approach provides a predictive distribution that quantifies the uncertainty of predictions, thereby enhancing the precision of MSI-H assessments and informing clinicians about the need for additional confirmatory lab testing. Additionally, by calculating the MSI-H status at both the tile and slide levels, our model provides new insights into the tumor microenvironment, as related to ICI responsiveness in gastric cancer.

## 2 Results

### Datasets

In this study, we analyzed H&E-stained WSIs from 12 distinct datasets comprised of colorectal and gastric cancers, with 2,091 and 1,101 slides, respectively. These datasets included a diverse patient population comprised of Asian, Black or African American, and White patients treated at multiple international sites, including Yonsei University and, Seoul St. Mary’s Hospital in Korea, Mayo Clinic in the USA, and various international sites (Table 1).

**Table 1:** Summary statistics of the data sets. For the GC-ICI cohort, we include 17 WSIs that have no MSI-H information, which were used in the analysis of the correlation between the predicted MSI-H region and immunotherapy response in Results section. W: White; B: Black or African American; As: Asian; AN: American Native or Alaskan Native; U: unknown.

| Cancer type | Dataset | Samples<br>N | MSI-High<br>N | Collection<br>sites | Race<br>N |
| --- | --- | --- | --- | --- | --- |
| Colorectal | TCGA-CRC | 361 | 65 (18%) | Various<br>international | W: 206 (57%)<br>B: 47 (13%)<br>As: 10 (3%)<br>AN: 1 (< 1%)<br>U: 97 (27%) |
|  | Yonsei-1 | 174 | 71 (41%) | Korea | As: 174 (100%) |
|  | Yonsei-1-remade | 146 | 53 (36%) | Korea | As: 146 (100%) |
|  | Yonsei-2 | 95 | 54 (57%) | Korea | As: 95 (100%) |
|  | STMary-Colon | 98 | 23 (23%) | Korea | As: 98 (100%) |
|  | CPATC-COAD | 221 | 53 (24%) | USA | W: 166 (75%)<br>B: 18 (8%)<br>As: 27 (12%)<br>AN: 3 (1%)<br>U: 7 (3%) |
|  | Mayo clinic | 966 | 255 (26%) | USA | W: 911 (91%)<br>B: 5 (< 1%)<br>As: 3 (< 1%)<br>AN: 10 (1%)<br>U: 67 (7%) |
| Stomach | TCGA-STAD | 284 | 60 (21%) | Various<br>international | W: 178 (63%)<br>B: 12 (4%)<br>As: 63 (22%)<br>U: 31 (11%) |
|  | Yonsei-Classic | 581 | 40 (7%) | Korea | As: 581 (100%) |
|  | STMary-GC | 72 | 22 (31%) | Korea | As: 72 (100%) |
|  | Molecular<br>Subtypes | 61 | 17 (28%) | Korea | As: 61 (100%) |
|  | GC-ICI | 103 | 14 (11%) | Korea | As: 103 (100%) |

The colorectal cancer datasets were TCGA-CRC, Yonsei-1, Yonsei-1-remade, Yonsei-2, STMary-Colon, CPATC-COAD, and Mayo Clinic. We used the TCGA-CRC and Yonsei-1 for training and the rest for validation. Of note, the Yonsei-1-remade dataset was generated by re-cutting slides and performing H&E staining from existing blocks of the Yonsei-1 dataset to explore how staining variability affected MSI-H prediction.

For gastric cancer, we analyzed datasets named TCGA-STAD, Yonsei-Classic, STMary-GC, GC-ICI, and Molecular sub-types. TCGA-STAD or Yonsei-Classic were used to train the model for gastric cancer sample analyses, and the remaining datasets were used for validation. The MSI status of the samples in the Molecular sub-types dataset was determined by both PCR and IHC. Thus, this dataset represents the gold standard for our validation efforts. Finally, the GC-ICI dataset consisted of gastric cancer patients treated with ICIs and allowed us to test the clinical utility of MSI-SEER to predict ICI response.

### Developing and Training the MSI-SEER model

The workflow of this study is summarized in Figure 1. Our tumor MSI status prediction model MSI-SEER consists of two main components (Figure 1-(A)). The first is a feature extractor that uses a pre-trained deep learning network to compute feature vectors from image tiles in a WSI within the transfer learning framework. The second is a prediction model based on a DGP in weakly supervised learning.

**Fig. 1:**
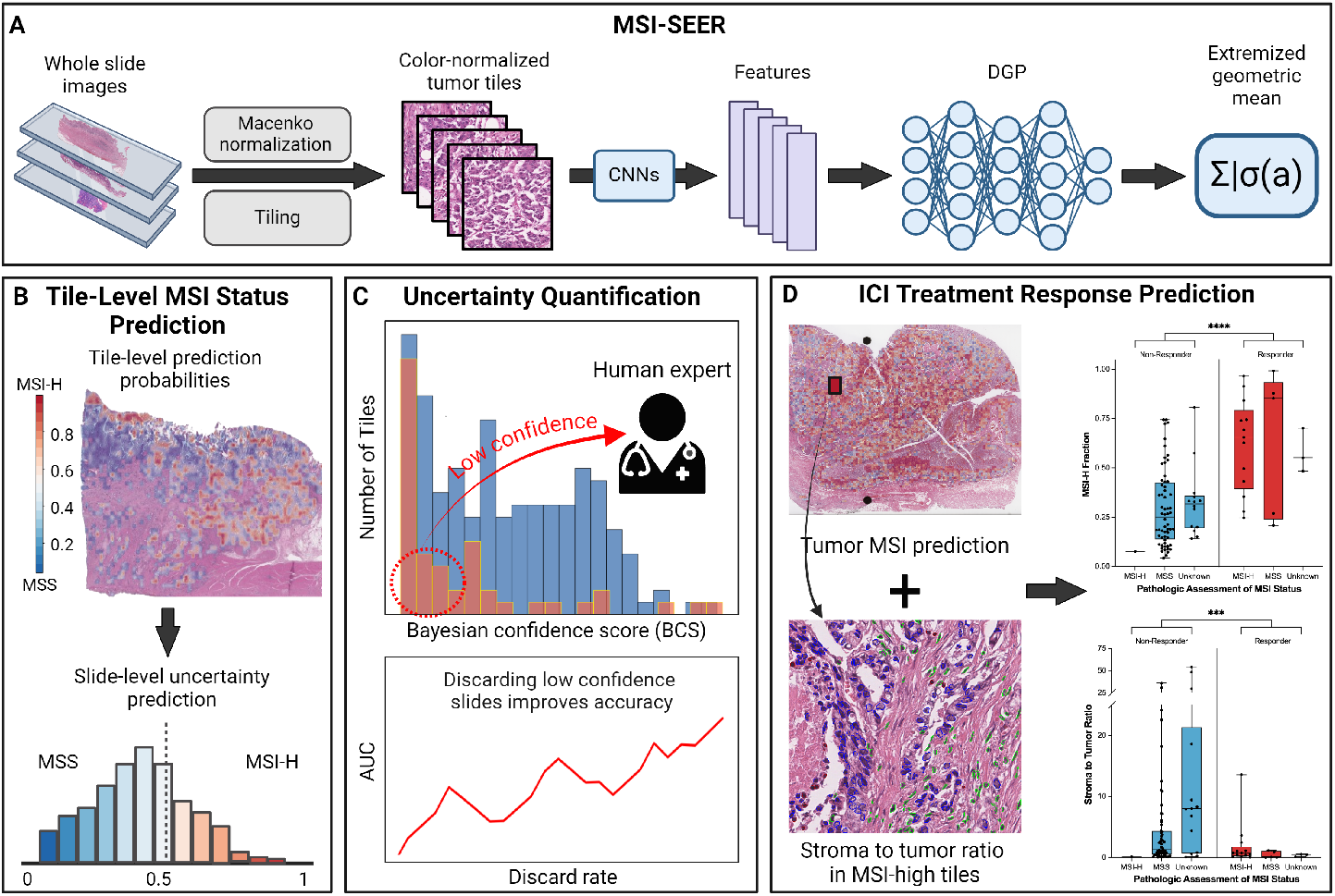
Workflow of summary of current study. (A) **MSI-SEER**. The method consists of two core components, an image feature extractor and a DGP-based MSI prediction model. Tumor tiles in a WSI were color normalized using Macenko method and transformed into feature vectors using a pre-trained CNN model. The slide-level MSI-H predictions were made by aggregating the tile-level MSI-H predictions using the weighted version of the extremized geometric mean of the tile-level MSI-H probabilities (*σ* is the sigmoid function and *a* is the extremized parameter). (B) **Output of MSI-SEER** provided MSI-H prediction probability at both the tile-level and slide-level and quantified predictive uncertainty at the slide-level. (C) **Uncertainty quantification** identified cases that should be referred to human experts for further investigation. Selective exclusion of highly uncertain predictions improved the model’s prediction performance. (D) Information provided by MSI-SEER predicted **immune check point inhibitor (ICI) treatment response**. We can predict ICI-treatment response using predicted MSI-H tumor region by **MSI-SEER** and stromal composition obtained by an image-based cell-type classification method CellViT.

#### Feature Extraction

Each WSI was divided into non-overlapping tiles, with each image set to 256 × 256 μm. Tiles containing mostly tumor were color normalized using the Macenko method [28] and were converted to image features by either CTransPath [29] or each of the nine MSIDETECT CNNs [12]. Then, these image features were inputted into our DGP models (see Methods for more details).

#### Model Inference

We first computed the probability of each individual tile being MSI-high using a DGP model, which was defined over the tiles in a WSI. The final prediction was made by aggregating individual tile predictions to obtain a slide-level MSI-H predictive probability. This summation was performed using the extremized geometric mean of the probabilities, which demonstrated superior calibration performance for aggregating multiple predictions among other pooling methods in terms of the Brier score [30]. Furthermore, we assigned a weight to each tile within the geometric mean of the odds pooling operator, thus minimizing attention to irrelevant tiles for MSI prediction.

#### Model Training

We trained the DGP models using dropout variational inference [31] since random feature expansion [32] a DGP can be reduced to a specific structure of Bayesian Neural Network (BNN) [33]. Furthermore, since each WSI may contain thousands of image tiles, we randomly selected up to 300 tiles in each slide to train the model, based on a recent study that the random subsampling outperforms other sampling approaches for multiple-instance binary classification using WSIs when enough tiles (100 to 1000) are selected [34].

We first explored the DGP models based on the CTransPath feature extractor to determine the optimal number of GP layers. We trained the model with different numbers of layers, from 1 to 7, and selected the best one based on the 3-fold cross validation (CV) performance (Supplementary Figure 1). While we did not see a wide variation in performance with different numbers of GP layers, the model performance has the increasing pattern up to six GP layers, which was then used for all the training datasets for both DGP models integrated with CTransPath and with MSIDETECT CNN models.

We implemented our model in ensemble learning, where the model was trained using bootstrapped samples of training data 10 times, and the final dropout samples from all ensemble models were aggregated to make the final prediction. Of note, the DGP integrated with MSIDETECT CNN models included nine different ensemble models depending on which CNN model was used for feature extraction. Since selecting the best performer from the nine ensemble models for a test slide is challenging, we combined these nine ensemble models by using the same aggregation method used to combine 10 models trained using bootstrapped samples. Finally, we observed that the DGP integrated with MSIDETECT CNN models generally outperformed the DGP integrated with CTransPath (Supplementary Table 1-2). Unlike the MSIDETECT CNNs, the CTransPath model was trained without MSI status labels in self-supervised learning. Therefore, we will utilize the DGP models integrated with MSIDETECT CNNs, which we will refer to as MSI-SEER.

In the model performance heatmaps (Figure 2 and 3), *best* and *worst* for MSI-SEEER represent the best and worst performing models among the nine DGP ensemble models. The aggregate MSI-SEER had performances that were comparable to the best performing individual MSI-SEER models in most cases.

**Fig. 2:**
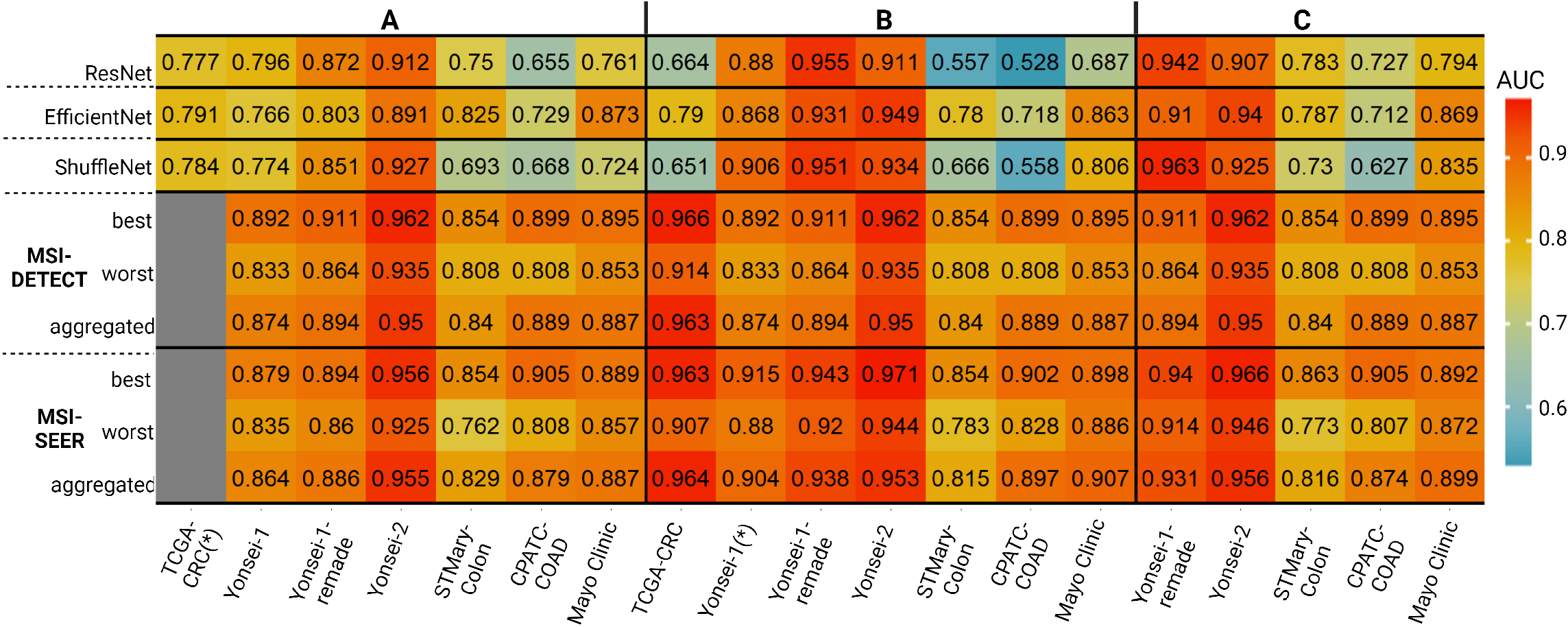
MSI-SEER performance for colorectal cancer. (*) denotes the dataset used for training and the remaining datasets were used for validation. Area under the ROC curve values are shown. The 3-fold cross-validation (CV) performance is evaluated for the training data, while the inter-cohort performance is evaluated for the validation datasets. MSI-SEER, ResNet, EfficientNet, and ShuffleNet were trained using (A) TCGA-CRC, (B) Yonsei-1, and (C) the combined data of TCGA-CRC and Yonsei-1. The 3-fold CV performance of MSIDETECT and MSI-SEER on TCGA-CRC was not evaluated because this dataset was already included in the training data for MSIDETECT.

**Fig. 3:**
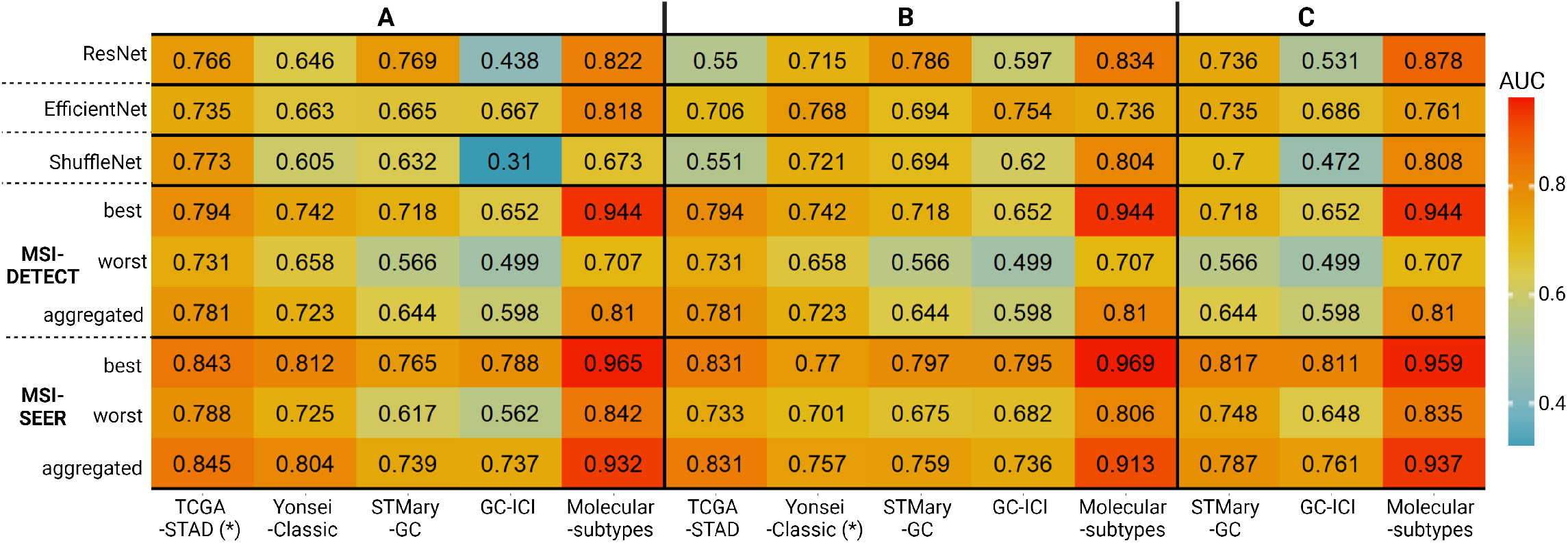
MSI-SEER performnace for gastric cancer. (*) denotes the dataset used for training and the remaining datasets were used for validation. Area under the ROC curve values are shown. The 3-fold cross-validation (CV) performance is evaluated for the training data, while the inter-cohort performance is evaluated for the validation datasets. MSI-SEER, ResNet, EfficientNet, and ShuffleNet were trained using (A) TCGA-STAD (B) Yonsei Classic and (C) the combined data of TCGA-STAD and Yonsei Classic.

For the colorectal datasets (Figure 2), we found that the MSI-SEER had better predictive performance for more cases when trained using Yonsei-1 than when trained using TCGA-CRC. However, we also found that the model trained on the combined data from both datasets did not perform better than the models trained on either dataset. This may be because the feature extractors were already trained on the datasets that included TCGA-CRC, and thus the combined data did not provide enough new information to MSI prediction. Therefor, for unseen colorectal cancer slides, we recommend using the aggregated MSI-SEER model trained on Yonsei-1. For the gastric cancer datasets (Figure 3), we found that the performance of the aggregated MSI-SEER was best when trained on a combined data from TCGA-STAD and Yonsei-Classic datasets, as compared to the models trained on each dataset alone. Thus, for unseen slides in gastric cancer, we recommend using the aggregated model trained on the combined TCGA-STAD and Yonsei-Classic datasets.

In the rest of the paper, MSI-SEER refers to the aggregated MSI-SEER model trained on Yonsei-1 for colorectal cancer and on the combined dataset for gastric cancer.

### MSI-SEER predicted MSI-status with accuracy similar to previously published models

To compare the predictive capability of MSI-SEER to previously published models, we adapted the experimental designs from Laleh et al. [35]. We compared MSI-SEER to previously reported CNN-based deep learning models, ResNet [36], ShuffleNet [37], and EfficientNet [38]. We also compared these models to the MSIDETECT CNN models. We evaluated the colorectal and gastric cancer samples separately so that each model was tested on the datasets only in the same cancer type to which the training dataset belonged. We did not observe a significant performance improvement when the models were trained using the datasets from both cancer types (Supplementary Table 3).

We trained ResNet, ShuffleNet, and EfficientNet using the same training steps as described by Laleh et al. [35], where the CNNs were trained in supervised learning under the assumption that all image tiles in a WSI shared the same label assigned to the WSI. We also attempted to retrain MSIDETECT, but observed severe performance degradation due to *catastrophic forgetting*, the phenomenon in which neural networks lose knowledge gained from previous tasks. We therefore used the pre-trained MSIDETECT models for the comparisons in the rest of the experiments. We used 3-fold CV to evaluate the performance of the training datasets (TCGA-CRC and Yonsei-1 for colorectal cancer and TCGA-STAD and Yonsei-Classic for gastric cancer). To evaluate the inter-cohort prediction performance, we trained each model using all data points in each training dataset, tested the models on the validation datasets in the same cancer type. Figures 2 and 3 show the prediction performance of the methods in terms of the area under the ROC curve (AUC) as a heatmap. We also used other metrics, recall, precision and F1 measure, to evaluate the performance of the models in Supplementary Tables 4 and 5.

In the model performance heatmaps, similar to MSI-SEER, *best* and *worst* for MSIDETECT represent the best and worst performing models among nine MSIDETECT CNNs. We observed that no single model among the nine MSIDETECT CNNs performed best for all datasets, and there was a wide variation between the best and worst performance (Supplementary Figure 2). Similar to MSI-SEER, we defined the aggregated model of the MSIDETECT CNNs by averaging the output MSI-H probabilities of the CNNs on a test WSI, where a slide-level prediction is also made by averaging the MSI-H probabilities over the tiles in the slide.

For colorectal cancer datasets (Figures 2), MSI-SEER had AUC ranging from 0.815 to 0.953, demonstrating that MSI-SEER worked well for colorectal cancer WSIs obtained from a diverse patient cohort. We next compared the MSI status prediction performance of the MSI-SEER model to the other models using DeLong’s method [39], which tests whether the AUC of one model is significantly different from that of another model. We found that MSI-SEER generally had comparable performance as the other methods, and in some cases, such as the CPATC-COAD and Mayo Clinic dataets, MSI-SEER showed better performance (Supplementary Figure 3). For the gastric cancer datasets (Figures 3), we first found that ResNet, ShuffleNet, and EfficientNet trained on TCGA-STAD, which consisted of a diverse patient cohort, performed worse on the validation datasets generated from Korean patients. However, their predictive performance did not improve much on these validation datasets even when the models were trained on the combined data from TCGA-STAD and Yonsei-Classic. In contrast, MSI-SEER performed well on these validation datasets generated from Korean patients, with AUC ranging from 0.761 to 0.937 (Figures 3C). Using DeLong’s method, we also showed that MSI-SEER significantly outperformed the other methods (Supplementary Figure 4).

### Incorporating predictive uncertainty improved MSI-SEER performance

For MSI-SEER, we quantified predictive uncertainty through a Bayesian confidence score (BCS) [40], where, 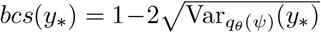, with 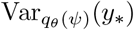 representing the variance of the slide-level predictive distribution. High BCS values indicated higher model confidence in the prediction. Our model’s inference, implemented using Monte Carlo dropout, estimated the likelihood of slide-level MSI-H status using extreme geometric means of tile log-odds. Higher computed variance occurred when the weighted sum of log-odds is near zero, indicating ambiguity in prediction (see Methods for more details). Unlike attention-based weakly supervised learning methods that aggregate tile-level features [29, 41], our method computed MSI-H probabilities for each tile and aggregated these to slide-level predictions. This tile-level analysis allowed for exploration of spatial MSI-H patterns within tumors, providing unique insights compared to other methods that may not offer detailed local predictions.

Figure 4 shows representative tile-level prediction results from our model, displaying the mean probability estimate of each tile being MSI-H on a heatmap on an WSI. Tiles with a dominant MSI-H or MSS morphological pattern generally yielded higher BCS, indicating more confident predictions (Figure 4-(A-B)). Conversely, tiles with heterogeneous MSI-H probabilities lead to lower BCS, reflecting higher uncertainty due to the complex patterns (Figure 4-(C-D)). In this case, despite the complex pattern, the weighted sum of the log-odds was approximately zero, and thus the slide-level MSI-H prediction probability was marginal (*≈*0.5) with the high uncertainty. WSIs with low predictive uncertainty (Figure 4-(C)) showed a spatially random distribution of MSI-H-like and MSS-like tiles. This correlation between spatial randomness and slide-level uncertainty was seen in each of our datasets, and there was a significant and strong negative correlation between the BCS and spatial (Altieri’s) entropy [42] (Supplementary Figure 5-6)

**Fig. 4:**
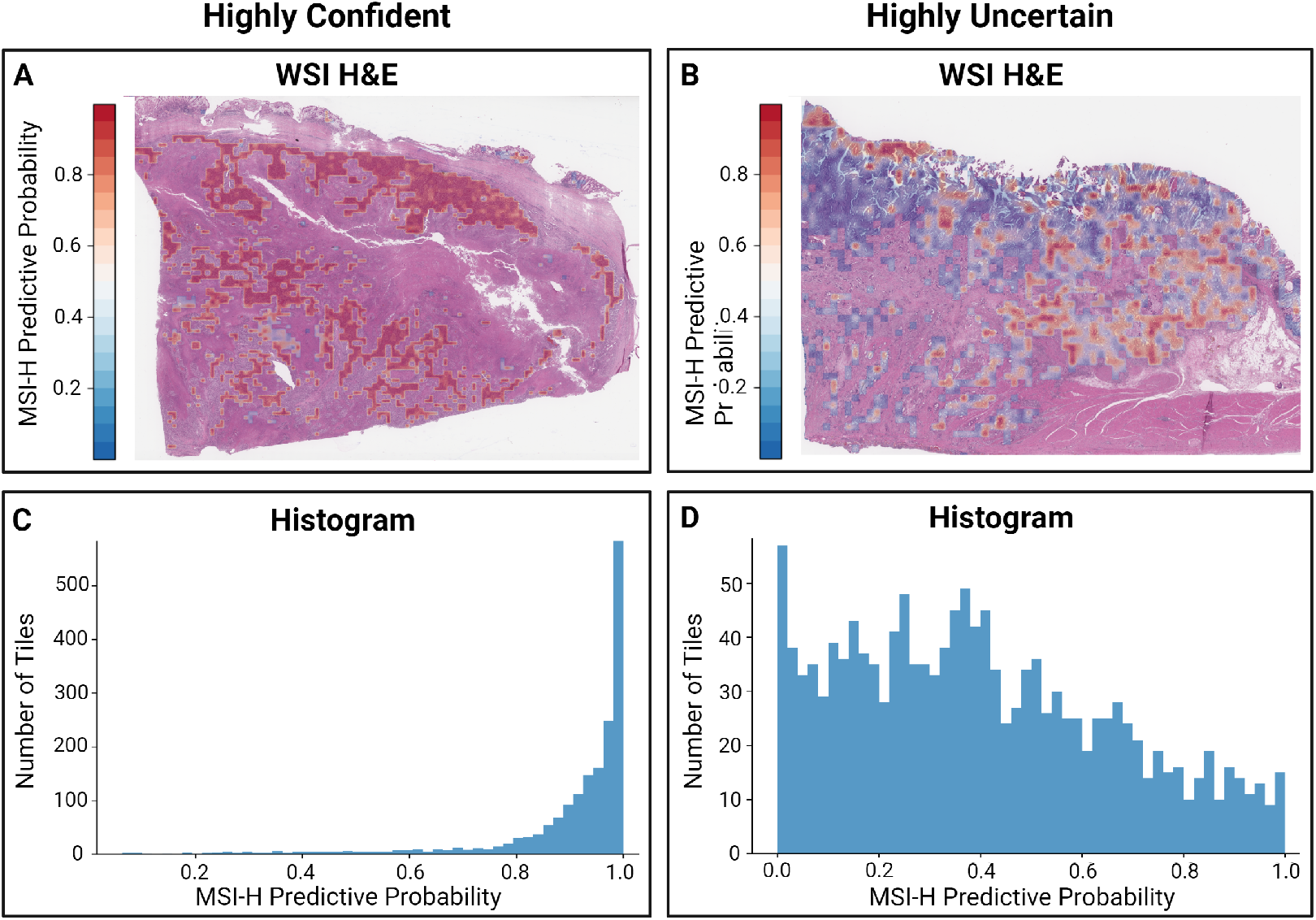
**Representative examples of uncertainty quantification in two gastric cancer samples**. (A-B) WSI images with tile-level MSI-H prediction probability heat maps (C-D) Histograms quantifying tiles based on predicted probability of MSI-H status. While both slides were considered MSI-H, the sum predictive probability was 0.996 (BCS=0.871) for the highly confident sample, and the sum predictive probability was 0.503 (BCS=1.6*e*^*−*5^) for the highly uncertain slide.

We next tested the effects of excluding the most uncertain predictions as based on Deodato et al [43]. We found that removing the most uncertain predictions enhanced the overall model performance by 1.1% and 2.6% in AUC in the colorectal (Figure 5A-B) and gastric cancer datasets (Figure 5C-D), respectively (Supplementary Table 6 for the other evaluation metrics). This selective approach of discarding the most uncertain predictions, consistently improved performance metrics, in contrast to random exclusions that showed no beneficial effect (Figure 3). These findings demonstrate the potential of leveraging predictive uncertainty to refine diagnostic accuracy, with further details on test data sets and various training scenarios show in Supplementary Figure 7-8.

**Fig. 5:**
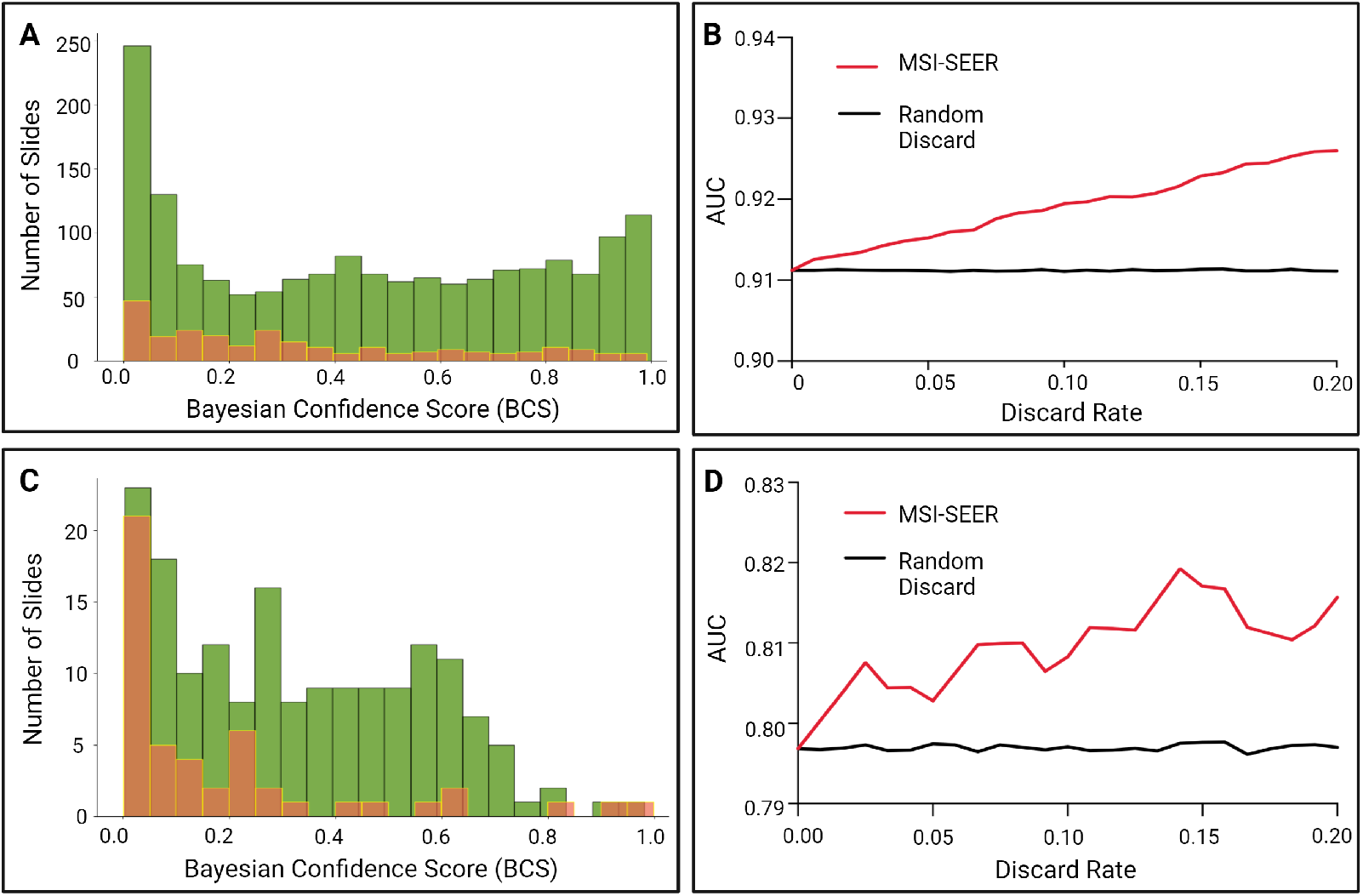
Examples of how to improve the predictive performance of our model using prediction uncertainty. In this model, the aggregated version of MSISEER was trained using Yonsei-1 in colorectal cancer or the combined data from TCGA-STAD and Yonsei-Classic in gastric cancer. (A) All test datasets in colorectal cancer, except the training data Yonsei-1, were combined and tested. The numbers of WSIs classified correctly are green, and those classified incorrectly are orange. The predictive uncertainty as measured by the Bayesian confidence scores are shown. (B) The changes in the prediction performance (in terms of area under the curve (AUC)) when the predictions are discarded at increasing rates, i.e. #discarded WSIs/#total WSIs, in each data cohort. The red line represents the change in the performance when the most uncertain predictions (as measured by the BCSs obtained by our model) are discarded while the black line is the average change in the performance where the predictions are randomly discarded 1,000 times at each rate. All the test datasets in gastric cancer, except the training data TCGA-STAD and Yonsei-1, were combined and tested. (C) The correctness of classification is shown. (D) The change in prediction performance is shown for the test datasets.

### Predicted MSI-H Regions and Stromal Composition Correlated to Immunotherapy Response in Gastric Cancer

To further assess the clinical utility of MSI-SEER, we performed a targeted analysis within the GC-ICI cohort, which included 21 slides from patients who responded to ICIs and 75 slides from patients who did not. We tested the association between the proportion of predicted tumor MSI-H regions and ICI response. To determine the MSI status for each tile, we used 0.5 as the cutoff for the mean predicted probability of the tile being MSI-H. We found that responders had an average of 62 % of tumor MSI-H predicted regions while non-responders had only an average of 30 % (*P <* 0.001, Figure 6A).

**Fig. 6:**
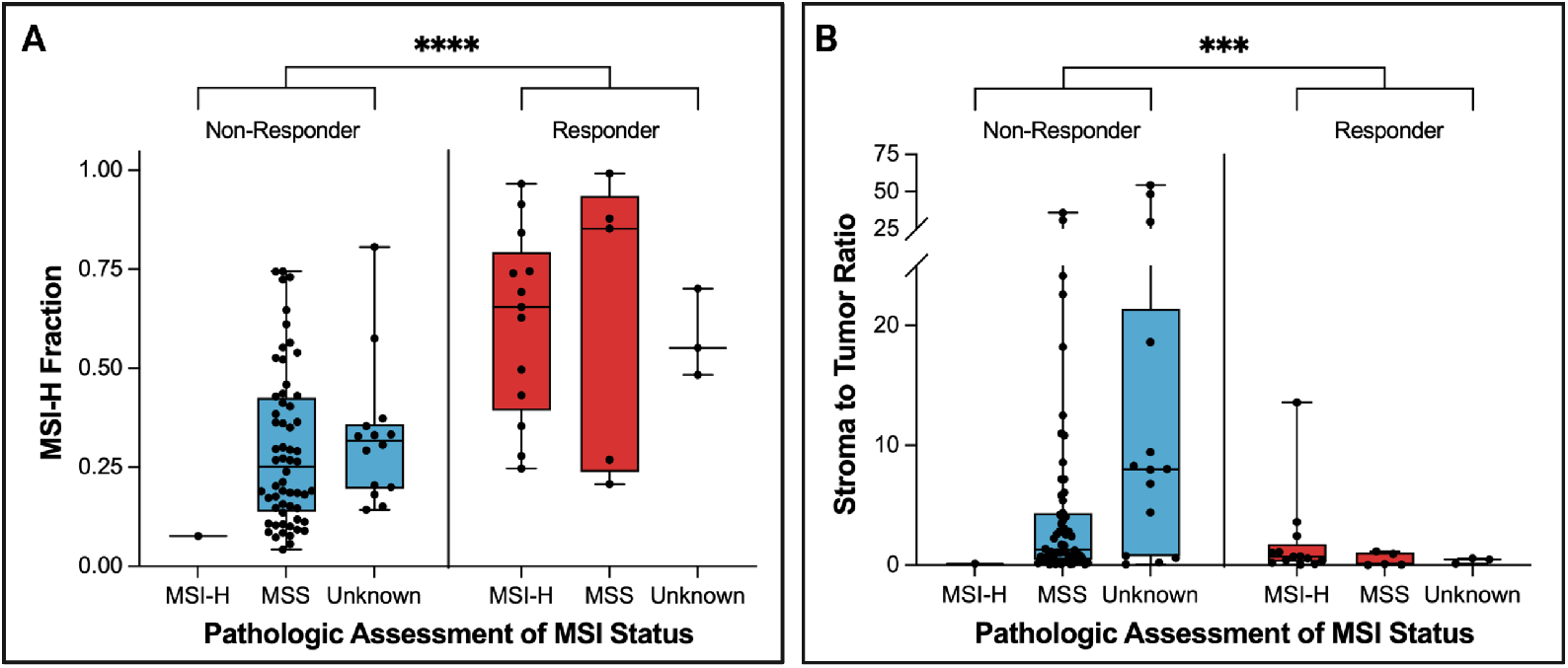
Comparison of MSI-H fractions in gastric cancer patients treated with immune checkpoint inhibitors (ICIs), stratified by treatment response (*N* = 96). (A) The MSI-H fraction, defined as the proportion of MSI-H tiles within a whole slide image (WSI). Responders demonstrated a significantly higher fraction of MSI-H tiles compared to non-responders (Wilcoxon test, *p* = 2.2*e*^*−*6^). To determine MSI-H and MSS tiles in a slide, we used 0.5 as the cutoff for the mean predictive probability of each tile being MSI-H over the dropout samples. MSI status is provided for reference and was not used in the comparison. Unknown denotes slides without available MSI status information. (B) Analysis of the stromal to tumor ratio within MSI-H predicted tiles, comparing responders and non-responders (Wilcoxon test, *p* = 0.0016). This metric assesses the microenvironment’s cellular composition, providing insights into the tumor-stroma dynamics that may influence ICI responsiveness.

Notably, there were 5 responders who were categorized as having MSS tumors by traditional testing methods. Three of the samples had more than 85.3% of tumor MSI-H predicted regions, and the other 2 had a similar amount of tumor MSI-H predicted regions as other confirmed MSI-H samples. Conversely, there was one clinically-determined MSI-H tumor that did not respond to ICI and it had only 7.7% of tumor MSI-H predicted regions. On review by board-certified pathologists (JHP and JYS), we found that these patients displayed histopathological features consistent with borderline state between MSS and MSI-H, suggesting that the MSI-SEER algorithm may uncover MSIH patterns not detected by standard laboratory tests. Thus, MSI-SEER may help refine patient selection for ICI therapy by identifying patients with MSS tumors who may benefit and MSI-H tumors who may not benefit.

Based on our previous work showing that stromal content is associated with ICI response [44], we next compared the stromal fraction within the predicted MSI-H tiles in responders and non-responders. Within each MSI-H tumor tile, we used CellViT to classify each cell as tumor or stromal [45]. Since the average number of tumor cells per tile was significantly higher in the responder group than in the non-responder group (Supplementary Figure 9), the stromal cell count was normalized by the tumor cell count. The majority of predicted MSIH tiles in non-responders contained a high number of stromal cells compared to those in responders. For example, more than 56% of the predicted MSI-H tiles in non-responders contain more than 50 stromal cells in the MSI-H tiles, while only 38.2% of the predicted MSI-H tiles in responders do. Figure 6 (B) shows that a high abundance of stromal cells in the MSI-H tiles is significantly associated with ICI non-responsiveness.

Finally, we developed a rule-based classifier incorporating predicted MSI-H fraction and stroma-to-tumor ratio within predicted MSI-H regions to predict ICI response. We treated 17 samples that do not have MSI status information in the cohort as test data and used the remaining samples to define the classification rule. Based on the data ranges of the predicted MSI-H fraction and the stroma-to-tumor ratio of responders obtained from samples that have MSI status information (Supplementary Figure 10), we defined a patient with an MSI-H fraction greater than 0.5 and a stroma-to-tumor ratio of less than 3.6 in a predicted MSI-H tile as a responder. Using this simple rule, we were able to stratify the test samples into responders and non-responders with an accuracy of 94.1% (Supplementary Table 7). These results demonstrate the potential of MSI-SEER to predict ICI responsiveness.

## 3 Discussion

Determining tumor MSI status provides clinically actionable information for both colorectal and gastric cancers. Deep learning models that analyze WSI may provide cost-efficient and widely accessible MSI testing. Indeed, previous work has shown that AI-based models have promising utility to predict MSI status for both colorectal and gastric cancers [14, 46, 47, 48] [10, 49, 50]. However, most of these models were not validated in diverse patient cohorts or did not report the racial makeup of their patient samples, thus raising the question of the generalizability of these models. The necessity of external validated in large and diverse patient cohorts have been demonstrated by several recent studies. Wagner et al. analyzed colorectal samples from multiple countries and found decreased generalizability of their model on samples from different races [47]. Similarly, Kather et al. found that their model for gastrointestinal cancers performed poorly on an Asian cohort when trained on data from predominantly non-Asian patients [10]. We were able to overcome these limitations with MSI-SEER, which is a prediction model based on DGP in weakly supervised learning, through extensive experiments on large datasets comprised of patients from diverse racial backgrounds and collected from multiple international sites. By training MSI-SEER on samples collected from diverse patient cohorts, we were able to improve our model performance.

Our Bayesian approach also provided significant advantages over traditional neural networks by quantifying uncertainty. Previous models are point estimation methods, which provide only binary (MSS or MSI-H) results. These outputs do not effectively capture or interpret uncertainty [51, 52], often leading to overconfident [53] and potentially misleading results especially in when used in complex decision-making processes such as clinical care. With MSISEER, we are able to provide uncertainty quantification while maintaining comparable prediction performance as previously developed deep learning MSI prediction tools, such as MSIDETECT. However, the ability of MSI-SEER to quantify the uncertainty in the prediction makes MSI-SEER a more clinically useful tool as uncertain results can be augmented with expert review for more nuanced decision making.

Finally, MSI-SEER may predict ICI response in gastric cancer. MSI-SEER was able to identify a subset of patients who were classified as MSS by traditional testing methodologies yet showed a clinical response to ICI treatment. This predictive capability extended to analyzing the stromal composition within MSI-H predicted regions. We previously found that high tumor ACTA2 expression, a marker of cancer-associated fibroblasts, is associated with ICI non-response [44]. With MSI-SEER, we similarly observed that increased stromal cells within tumors correlated with ICI non-responsiveness. By combining MSI-H fraction with stroma-to-tumor ratio we were able to achieve high accuracy in predicting ICI responsiveness in patients with gastric cancer. These findings highlight the significant influence of tumor microenvironment on therapy effectiveness.

To our knowledge MSI-SEER is the first computational model that predicts ICI response from WSIs in gastric cancers. Given that MSI-H gastric cancer may not be responsive to cytotoxic chemotherapies [3], using MSI-SEER to predict ICI response may spare select patients the toxicities associated with chemotherapy and allow them to receive more optimal treatment in the form of ICIs.

## 4 Acknowledgments

We acknowledge funding from the National Cancer Institute (NCI) with grant 1R01CA276690 and from the Department of Defense (DOD) under grant CA190578 awarded to THH. THH is further supported by the Eric and Wendy Schmidt Foundation’s AI Innovation Award through the Mayo Clinic Foundation. MFP is supported by the American College of Surgeons Resident Research Scholarship and the National Cancer Institute (3R37CA265967-01A1S1). SCW is a UT Southwestern Disease Oriented Clinical Scholar and supported by the National Cancer Institute (R37CA265967, R01CA276690, U54CA233306, and U54CA283766). This study was approved by Gangnam Severance Hospital Institutional Review Board (Approval No: 3-2021-0367). SHL is supported by a grant from the National Research Foundation of Korea (NRF-2022R1A2C2010644). The Mayo Clinic H&E images used in this manuscript are part of the Colon Cancer Family Registry (CCFR, www.coloncfr.org), which is supported in part by funding from the National Cancer Institute (NCI), National Institutes of Health (NIH) (award U01 CA167551).

## 5 Methods

### Datasets

The datasets used in this study are H&E stained colorectal and gastric cancer slides collected from multiple institutions containing multiple racial groups. For the colorectal cancer data, we first used a publicly available multi-center data from The Cancer Genome Atlas (TCGA) project: TCGA-CRC. Yonsei-1 and Yonsei-2 were collected from Gangnam Severance Hospital, Yonsei University College of Medicine, Seoul, Republic of Korea. Yonsei-1-remade was reprocessed from Yonsei-1: the remaining tumor tissues from a subset of the patients in Yonsei-1 were scanned to produce whole slide images. STMary-Colon was collected from Seoul St. Mary’s Hospital, Seoul, Republic of Korea. The CPATC-Colon dataset is from The Clinical Proteomic Tumor Analysis Consortium Colon Adenocarcinoma Collection (CPTAC-COAD). Mayo Clinic slides are from Colon Cancer Family Registry (CCFR).

For gastric cancer data, we also used publicly available data from TCGA, (TCGA-STAD). We then included Yonsei-Classic data which was collected from patients who received D1 gastrectomy plus capecitabine and oxaliplatin chemotherapy or surgery alone. Molecular sub-types dataset is from Seoul St. Mary hospital in Korea, and the GC-ICI dataset was obtained from Yonsei University College of Medicine and Seoul St. Mary hospital.

Note that TCGA (TCGA-CRC and TCGA-STAD), CPATC-colon and Mayo clinic datasets data contained primarily white patients, whereas all other datasets were collected from Asian (Korean) patients. Table 1 contains the summary statistics of each dataset in both colorectal- and stomach cancers. Note that images without clinical MSI status were excluded from experiments where MSI-H prediction performance was evaluated.

#### Related Work: Weakly Supervised Learning and Prediction Uncertainty Analysis

Prediction tasks using WSIs can naturally be formulated as a weakly supervised learning problem or a set problem [54]. Typical WSIs can exceed gigapixels in size, so one of the most common approaches is to divide a WSI into multiple, non-overlapping small image tiles. In contrast to standard supervised learning, where an input point and corresponding label are paired, in the weakly supervised learning that we utilized for the current study, a set of input points (image tiles) are given a slide-level single label, which in our study is the PCR-determined MSI status or IHC-detection of mismatch repair protein presence. Multiple instance learning (MIL) [55], a special case of weakly supervised learning, is defined in a similar way with a bag (a slide) of instances (image tiles in a slide). A label is available for a bag, not for individual instances in a bag. Despite its application to WSIs for tumor detection and MSI status prediction, MIL may not align perfectly with our task due to the restrictive assumptions regarding the label generation process. Since MIL assumes that the label of a bag is determined by the presence of positive instances [55], even if there are only a few MSI-H-like tiles in a slide and all other tiles are non-MSI-H, the label of the slide is set to MSI-H, which might be too restrictive, due to intratumor heterogeneity. For example, a WSI of a MSS tumor must be allowed to include a few MSI-H like image tiles, if those images are present. We therefore use the term ”weakly supervised learning” rather than ”multiple instance learning” to describe the MSI prediction problem. In practice, many early MSI prediction methods [10, 12, 14, 16, 17] do not fully solve the prediction problem in weakly supervised learning, due to simplicity in implementation and computational limitations. These models were trained in standard supervised learning, which trains models to associate each image tile with its slide-level label. Within the scope of weakly supervised learning, the prediction process was restricted to the aggregation of individual tile predictions within the slice by mean or max pooling. However, due to intratumor heterogeneity, some image tiles, e.g., MSI-H-like tiles in an MSS slide or MSS-like tiles in an MSI-H slide, may introduce noise into the models, and all tiles in a slide may not be relevant for prediction. To address these issues, we implement both training and prediction in weakly supervised learning, where the probability of a slide being MSI-H is computed by aggregating the probability estimates from the image tiles in the slide by the (extremized) geometric mean of odds [30]. In addition, the weighted extension of the geometric mean of odds can help the model to disregard irrelevant tiles in the prediction.

Quantifying the uncertainty in the prediction allows us to understand what a prediction model does or does not know. A high degree of uncertainty at a specific test point may indicate that the model’s lack of confidence in its prediction, possibly because the test point is out of the training data distribution, or because there are unknown variables or noise within the observations. These underlying causes of high prediction uncertainty align closely with two different types of uncertainty: epistemic and aleatoric uncertainty [56, 57]. Epistemic uncertainty is related to the randomness of the model parameters due to the insufficient number of training data and can be reduced if we collect more data. Conversely, aleatoric uncertainty refers to intrinsic noise in the observations and is irreducible. Once the prediction uncertainty is computed, we can refer the test points, or WSIs in this context, with high uncertainty to human experts for in-depth evaluation. This step can potentially reduce prediction errors on challenging test points, thereby enhancing the overall performance of the prediction model. Standard deep learning-based methods, limited to providing point estimates, are unable to capture prediction uncertainty [51]. The final outputs of these methods (e.g., softmax probabilities in neural networks) are frequently misinterpreted as uncertainty, but unfortunately, they are known to be overconfident for test points far from the training data [53] and miscalibrated [52]. On the other hand, Bayesian approaches can intrinsically generate uncertainty in the prediction, providing a distribution over a prediction in Bayesian learning (by averaging the likelihood over the posterior distribution of the model parameters). Gaussian Process (GP) [58], a nonparametric Bayesian method for nonlinear function estimation, allows us to compute a prediction distribution in the form of a Gaussian distribution, where the variance captures the uncertainty in the prediction. DGP [26, 59], a multi-layer hierarchical extension of GPs, inherits the attractive properties of GPs, including nonparametric prior modeling and well-calibrated uncertainty estimation, while providing a more flexible and generalizable prior distribution than GPs. It is noteworthy that a GP is nothing more than a special case of a DGP (i.e., a single-layer DGP).

### Image preprocessing

Each pathology image was divided into multiple non-overlapping patches (the size of each image tile is set to 256 × 256 *μ*m). Only image tiles containing mainly tumor were used in the experiments: an image tile containing non-tissue regions or consisting of *≥*50% of white background was discarded. To detect tumor from image patches, we trained a ResNet-18 model on TCGA-CRC data. All remaining image tiles were color normalized using Macenko normalization method [28] to suppress possible variations across samples or different data cohorts. Then, each image patch was fed to the trained tumor detection model.

### Feature transfer learning

For our DGP model, we used transfer learning to extract features from image patches: a feature vector for an image tile was calculated using a pre-trained model (feature extraction in transfer learning [60]).

### Problem definition

The task of predicting the MSI status from a whole slide image can be defined in weakly supervised learning. We are given pairs of an input image and an output label, i.e., 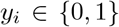, where *ℐ*_*i*_ represents the *i*th image, *y*_*i*_ is a binary label, i.e., *y*_*i*_ *∈ {*0, 1}, where *y*_*i*_ = 1 for MSI High and *y*_*i*_ = 0 for MSS, and *N* is the total number of the training images. Each image is divided into multiple non-overlapping small image patches each of which can be processed separately. Using transfer learning to deal with a small number of training labels, each image *ℐ*_*i*_ can be represented by a set of image feature vectors, i.e., 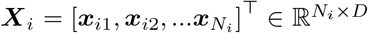, where ***X***_*i*_ *∈*ℝ^*D*^ and *N*_*i*_ is the number of image patches in the *i*th image. All training input data can be denoted by 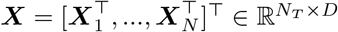, where 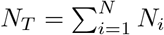, and all training output labels by ***y*** = [*y*_1_, …, *y*_*N*_]^⊺^. The objective is to learn a classifier that takes a set of *N* image patches of a new image, i.e., 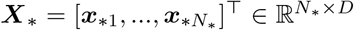 as input and estimates its prediction probability given the training data i.e., *p*(*y*_*\**_ = 1 | ***X***_*\**_, ***X, y***).

In order to deal with the weakly supervised learning problem described above, where there is only a single slide-level label available for a set of multiple feature vectors computed from image tiles in a WSI, we propose to use the aggregation of image tile-level probability estimates of MSI-High based on the geometric mean of the odds operator [30]. We first assume that we can access the probability of each images tile (***x***_*ij*_) being MSI-High in the *i*th WSI, i.e., *p*_*ij*_ = *P* (*y*_*ij*_ = 1 | ***x***_*ij*_) and that the log-odds (of the tile-level probability estimates) are sampled from the Normal distribution centered at the true log-odds as in [30]. Then, the maximum likelihood estimator of the true log-odds in this model setting is given by the geometric mean of the odds:

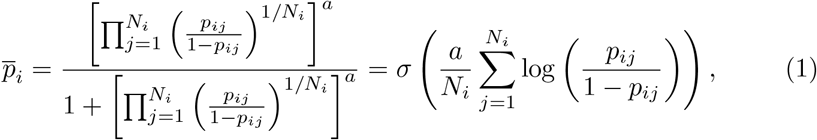

where *σ* is the sigmoid function and *a >* 0 is the extremization parameter. Note that a larger value of *a* makes 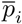 more extreme (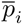 becomes closer to either 1 or 0). We then introduce a simple extension of (1) to the uneven tile weights by replacing the uniform weights (1*/N*_*i*_) with weight terms 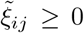 (with 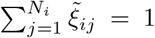). We define a nonlinear function mapping from image tile vectors, i.e., ℝ ^*D*^, to ℝ ^2^ each output dimension of which are denoted by [*ϕ*]_1_ or [*ϕ*]_2_: [*Φ*]_1_ directly models the log-odds of each image tile, i.e., 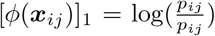 and [*ϕ*]_2_ models the weights with additional transformations, i.e., 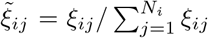, where *ξ*_*ij*_ = 0.1 + 0.9 *\* σ*([*Φ*(***x***_*ij*_)]_2_). To ensure notational consistency throughout the paper, we define the (normalized) likelihood of the mapping functions *ϕ* with the introduction of a loss function *l* as follows [61]:

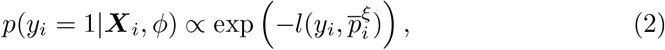

where 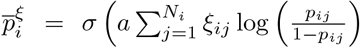 is the uneven weighted extension of (1). Note that, when the cross entropy is used as the loss function, i.e., 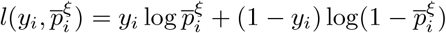, where the right side on (2) becomes nothing but 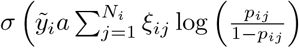, where 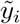 is the signed binary label, i.e., 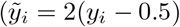.

#### Deep Gaussian processes (DGPs) with random feature expansion

This subsection shows that the mapping function *Φ* can be modeled using a DPG with random feature (RF) expansions [32, 33]. More formally, we assume that *ϕ* is modeled by *L* layers of GPs (i.e., a DGP with *L* layers):

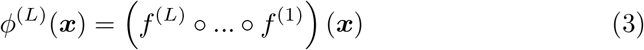

where the superscript (*l*), where 1 ≤ *l* ≤ *L*, denotes the *l*th layer and *f* ^(*l*)^ in each layer is a multivariate function whose the output dimensionality is *D*^(*l*)^, i.e., 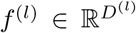. Each output dimension is assumed to be modeled by an individual GP, and thus there are *D*^(*l*)^ number of GPs in each layer.

To understand this modeling more clearly, let us consider the latent function values of the all the training data points ***X*** up to the *l*th layer: 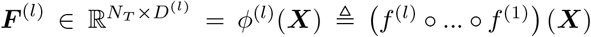. Assuming that all the GPs in each layer share the same covariance matrix, we have 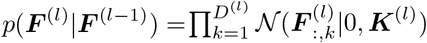, i.e., each column 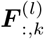 is modeled by a GP with the covaraince matrix 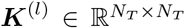 whose (*i, i*^*′*^) element is defined over the output function values of the previous layer i.e., 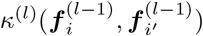, where *κ*^(*l*)^ is the covariance function the *l*th layer and ***f*** _*i*_ is the *i*th row vector of 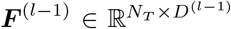, i.e., 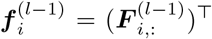. Note that, the layer depth at zero is defined to be the input layer, i.e., 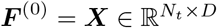.

Note that, the total number of instances, *N*_*T*_, can be massive even if the number of images *N* is small (e.g., the number of whole slide images is a few hundreds, but each image can have thousand image tiles). In addition, the memory space of and the time complexity of algebraic operations on each covariance matrix are 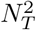 and 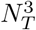, respectively, which makes a GP prohibitive even for a dataset of hundreds of images. To alleviate this computational issue, we consider the low-rank approximation of the covariance matrices *{****K***^(*l*)^*}*:

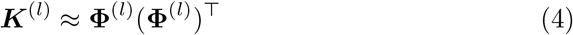

where 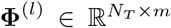 and *m* ⩽ *N*_*T*_. This approximation leads a Bayesian linear model that can approximate the GP latent function values [62]. Using the notational abuse, let us define ***F*** ^(*l*)^ ≜ **Φ**^(*l*)^***W*** ^(*l*)^, where the priors over the linear weight matrix 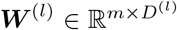 assumed to be i.i.d. Gaussians, i.e., 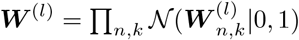. One can easily see the validity of this linear model approximation by checking cov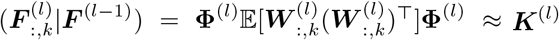.

To implement the low-rank approximation in (4), we employ random feature expansions [32, 33]. First consider the arc-cosine kernel function as the covariance between two input points ***h*** and ***h***^***′***^ in [63] (in our case the input ***h*** is assumed to be the output of the previous layer of the DGP model, i.e., 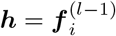 and 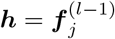 with arbitrary indices *i* and *j*):

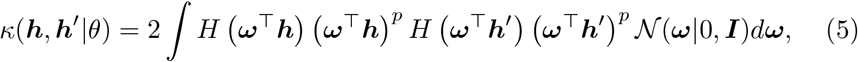

where *H* is the Heaviside step function. Note that we can approximate the integration with finite samples drawn from the Gaussian (*ω*_1_, …, *ω*_*m*_ *∼* 𝒩 (***ω*** |0, ***I***)). With the fact that when 1, *H*(*·*)(*·*)^*p*^ becomes ReLU(*·*), the low rank matrix **Φ**^(*l*)^ in the approximation (5) can be written by

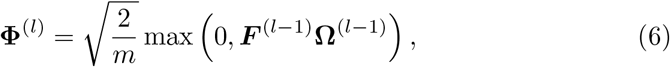

where 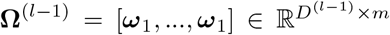 and 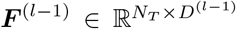 are the outputs of the previous layer by the definition. Recalling that ***F*** ^(*l*)^ = **Φ**^(*l*)^***W*** ^(*l*)^ and ***W*** ^(*l*)^ or **Ω**^(*l*)^ can be defined as network weights with a prior distribution (Gaussian), a DGP with random feature expansion can be reduced to a Bayesian deep NN.

### Model inference

We train the model in Bayesian learning framework with the black-box (BB) *α*-divergence minimization [64, 65] which can be, roughly speaking, understood as a stochastic gradient version of power expectation and propagation (EP) [66]. Power EP, which is based on the local *α*-divergence minimization [67], generalizes EP to include variational inference (VI) (*α→* 0) or EP (*α* = 1) as a special case with a general setting of the parameter *α*. However, it does not scale well because its implementation involves storing a local approximation parameter (also known as a site parameter) of each likelihood factor (each data point) in memory. Furthermore, since Power EP (as well as EP) is based on message passing, its solution is not guaranteed to converge to a stationary point of the energy function. On the other hand, since the BB-*α* divergence minimization directly optimizes the energy function with respect to the global (single) parameter which is combined from the site parameters without performing message passing [64], we can directly apply any gradient descent methods or desirably stochastic gradients methods for the optimization and thus this method is applicable for large-scale problems. In particular, in this work we use the further approximate version of BB-*α* divergence minimization proposed in [65] because it leads a simple and efficient (variational) inference method along with the use of Monte Carlo Dropout [31, 68].

We first define the approximate posterior distribution over the model parameters 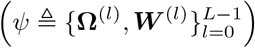. We show here only the case of the linear model parameter ***W*** ^(*l*)^ (the superscript for the layer number is omitted for rotational brevity), but the posterior distributions over the other variables can be defined in exactly the same way. The posterior distribution over ***W*** is defined as a mixture of two Gaussian distributions [31]:

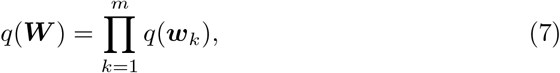

where

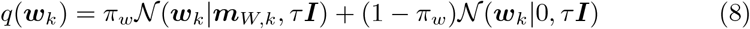

where *π*_*w*_ ∈ [0, 1]. Let us then define ***M*** _*W*_ = [***m***_*W*,1_, …, ***m***_*W,m*_]. Introducing a binary variable vector ***z***_*w*_ each element of which follows the Bernoulli distribution, i.e., ***z***_*w,k*_ *∼* Bernoulli(*π*_*w*_) for *k* = 1, …, *m* and letting *τ* tend to zero, random samples drawn from the posterior (7) can be approximated by [31]

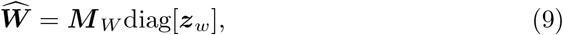

where the operator diag[***v***] is a diagonal matrix with the vector ***v*** and each element in the binary vector ***z***_*w*_ can switch on or off the corresponding column of 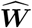 with the probability *π*_*w*_. We apply the same construction to the posterior distributions of the other variables and define the variational parameters 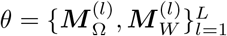. Now, we can define the BB-*α* divergence energy function [65] of our model for a mini-batch 𝒪_*b*_ using Monte Carlo expectation with random samples 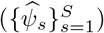 drawn from the posterior distribution over the model parameters which is parameterized by the variational parameters *θ*, i.e., *q*_*θ*_(*ψ*):

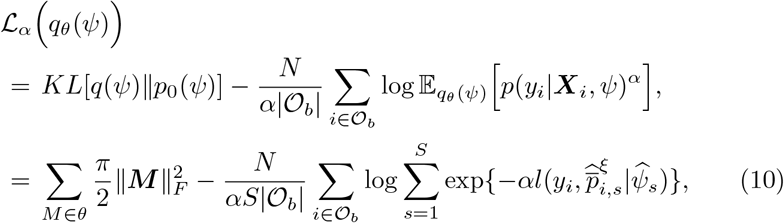

where *π* = *π*_*w*_ = *π*_Ω_ and 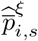 is a forward pass of 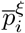at the parameter 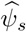, *p*_0_ is the prior distribution (a Gaussian distribution for one column in each parameter matrix) and the Kullback-Leibler (KL) divergence in the first line was also approximated as in [31]. To generate a mini-batch 𝒪_*b*_, we randomly draw the same number of samples from the positive and negative classes.

### Prediction for a new test image

The predictive distribution of a test image (feature vectors) ***X***_*\**_ can be computed as

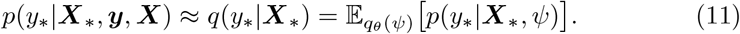

Again, the above expectation can be approximated using random samples of the model parameters by resampling the binary Bernoulli variables (i.e., using Monte Carlo dropout). For example, the predictive distribution of a test image being MSI-H is given by 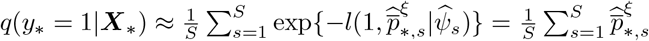 (recall that *σ* is the sigmoid function and 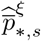 is a forward pass of 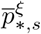 from the DGP model with the sampled model parameter 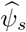 at the test image ***X***_*\**_). To evaluate the uncertainty in prediction, we calculate the variance of the predictive distribution as follows [69]

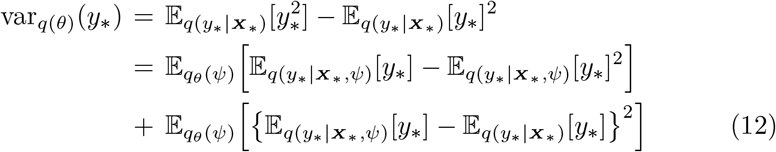

where the first term in the last equation is aleatoric uncertainty and the second term epistemic uncertainty [57, 69]. Again, the variance can be approximated with random samples drawn from *q*_*θ*_(*ψ*):

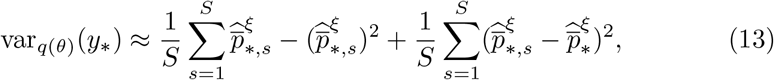

where 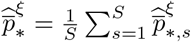.

### Performance evaluation

For the evaluation metrics, we basically used the area under the ROC curve (AUC), whose value range is from 0 to 1. An AUC close to 1 indicates that a model has good predictive power. To compare the performance of two classification methods, we used DeLong’s method [39], which tests whether the AUC of one model is significantly different from that of another model.

