## supplementary information for "Deep Gaussian Process with Uncertainty Estimation for Microsatellite Instability and Immunotherapy Response Prediction Based on Histology"

#### Supplementary Methods

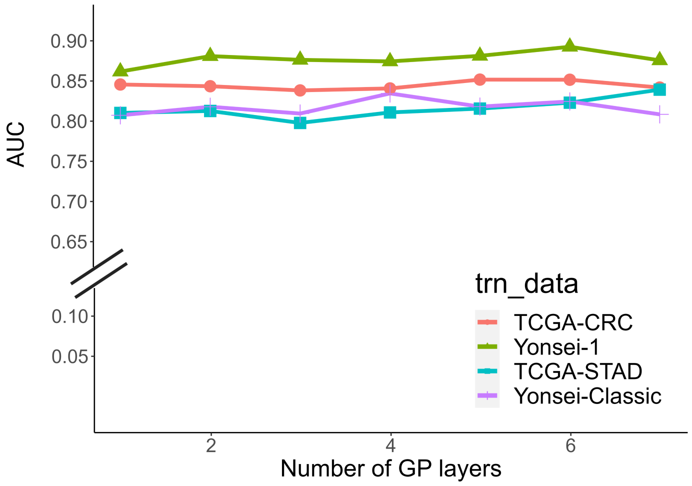


**Supplementary Figure 1**: The 3-fold cross validation prediction performance of our model (DGP+CTransPath) on the training datasets (TCGA-CRC, Yonsei-1, TCGA-STAD, Yonsei-Classic) at different numbers of Gaussian processes (GP) layers.

Our DGP model has several user-defined hyperparameters, including the value of $\alpha$ in the black-box $\alpha$ divergence formulation (eq. 10 in Methods), the rank of the approximate covariance matrix in each GP layer ($m$), and the number of layers in a DGP model ($L$). For the black-box $\alpha$ divergence formulation, it reduces to expectation and propagation (EP) when $\alpha = 1$ or to variational inference (VI) when $\alpha$ converges to 0. We specifically set $\alpha = 0.5$ for all the experiments because it has been reported that the black-box $\alpha$ divergence algorithm with the non-standard setting of $\alpha$, e.g., $\alpha = 0.5$, outperformed EP ($\alpha = 1$) or VI ($\alpha \to 0$) [1]. For the rank ($m$), we used the same fixed value ($m = 100$) for all GP layers in all the experiments.

To determine the optimal number of GP layers in the DGP models, we evaluated the prediction performance of the DGP model integrated with CTransPath (DGP+CTransPath) by varying the number of GP layers from 1 to 7. The prediction performance was evaluated by AUC on the training datasets in the main text. We found that the performance did not vary significantly with the number of GP layers (Supplementary Figure 1). For simplicity, we kept the number of GP layers fixed ($L = 6$) for all the experiments in the study,

To train our DGP models, i.e., to optimize the objective function (eq. 10 in Methods), we use stochastic gradient methods. Specifically, we used Adam optimizer with $l = 0.001$ (the learning rate), $\beta_1 = 0.9$, $\beta_2 = 0.999$ and $\epsilon = 1e^{-7}$. The maximum number of epochs used for model training was set to 100.

We trained CNN-based deep learning models according to the same training steps described by Laleh et al [2] for comparison. We used ResNet, ShuffleNet, and EfficientNet as the backbone CNN Models pretrained on ImageNet. The models were finetuned end-to-end for the MSI prediction task with a training epoch of 8 and a patience of 5. Training was stopped if the validation loss did not decrease. For all CNNs, we trained models with learning rates set to 1e-4, weight decay at 1e-5, batch size of 512; using Adam optimizer and freeze ratio of layers at 0.5. Both DGP models and CNNs were implemented in PyTorch using Python 3.7.

### Supplementary results

In this section, we provide more detailed results for the experiments included in the main text and perform additional experiments.

| Feature extractor | TCGA-CRC | Yonsei-1 | Yonsei-1 -remade | Yonsei-2 | STMary Colon | CPATC COAD | Mayo Clinic |
|---|---|---|---|---|---|---|---|
| MSIDETECT | Training Set | 0.864 | 0.886 | 0.955 | 0.829 | 0.879 | 0.887 |
| CTransPath | | 0.831 | 0.859 | 0.932 | 0.824 | 0.860 | 0.879 |
| MSIDETECT | 0.964 | Training Set | 0.938 | 0.953 | 0.815 | 0.897 | 0.907 |
| CTransPath | 0.805 | | 0.964 | 0.923 | 0.769 | 0.789 | 0.838 |
| MSIDETECT | Combined as Training Set | | 0.931 | 0.956 | 0.816 | 0.874 | 0.899 |
| CTransPath | | | 0.939 | 0.924 | 0.812 | 0.820 | 0.881 |

**Supplementary Table 1**: Comparison of MSI prediction performance in colorectal cancer of DGP models integrated with either the MSIDETECT or CTransPath feature extractors. When the MSIDETECT CNNs were used as the feature extractor, we consider only the aggregate model. AUC values are presented here.

| Feature extractor | TCGA-STAD | Yonsei-Classic | STMary-GC | GC-ICI | Molercular subtypes |
|---|---|---|---|---|---|
| MSIDETECT | Training Set | 0.804 | 0.739 | 0.737 | 0.932 |
| CTransPath | | 0.744 | 0.688 | 0.766 | 0.671 |
| MSIDETECT | 0.831 | Training Set | 0.759 | 0.736 | 0.913 |
| CTransPath | 0.795 | | 0.706 | 0.801 | 0.914 |
| MSIDETECT | Combined as Training Set | | 0.787 | 0.761 | 0.937 |
| CTransPath | | | 0.707 | 0.778 | 0.855 |

**Supplementary Table 2**: Supplementary Table 2: Comparison of MSI prediction performance in gastric cancer of DGP models integrated with either the MSIDETECT or CTransPath feature extractors. When the MSIDETECT CNNs were used as the feature extractor, we consider only the aggregate model. AUC values are presented here.

| Model | Yonsei-1-re | Yonsei-2 | STMary-colon | CPATC | Mayo clinic | ST-Mary GC | GC-ICI | Molecular sub-types |
|---|---|---|---|---|---|---|---|---|
| ResNet | 946 | 0.935 | 0.681 | 0.772 | 0.827 | 0.664 | 0.350 | 0.942 |
| EfficientNet | 0.904 | 0.928 | 0.554 | 0.756 | 0.777 | 0.773 | 0.526 | 0.846 |
| ShufflNet | 0.946 | 0.927 | 0.568 | 0.656 | 0.778 | 0.681 | 0.295 | 0.804 |
| DGP+CTransPath | 0.934 | 0.967 | 0.826 | 0.817 | 0.815 | 0.778 | 0.791 | 0.983 |
| MSI-SEER (best) | 0.940 | 0.976 | 0.850 | 0.877 | 0.863 | 0.855 | 0.818 | 0.945 |
| (worst) | 0.919 | 0.931 | 0.770 | 0.778 | 0.801 | 0.755 | 0.633 | 0.809 |
| (aggr) | 0.938 | 0.960 | 0.822 | 0.874 | 0.856 | 0.825 | 0.769 | 0.911 |

**Supplementary Table 3**: MSI prediction performance of the models trained using combined data from both colorectal and gastric cancer datasets (TCGA-CRC, Yonsei-1, TCGA-STAD, and Yonsei-classic). AUC values are presented here.

| Model | TCGA-CRC (precision, recall, F1) | Yonsei-1-re (precision, recall, F1) | Yonsei-2 (precision, recall, F1) | STMary-Colon (precision, recall, F1) | CPATC-COAD (precision, recall, F1) | Mayo Clinic (precision, recall, F1) |
|---|---|---|---|---|---|---|
| MSI-SEER | (0.839, 0.800, 0.819) | (0.825, 0.887, 0.855) | (0.962, 0.944, 0.953) | (0.778, 0.609, 0.683) | (0.761, 0.660, 0.707) | (0.709, 0.820, 0.760) |
| DGP+CTransPath | (0.440, 0.677, 0.533) | (0.906, 0.906, 0.906) | (0.959, 0.870, 0.913) | (0.959, 0.870, 0.913) | (0.500, 0.792, 0.613) | (0.549, 0.722, 0.624) |
| ResNet | (0.820, 1.000, 0.901) | (0.926, 0.946, 0.936) | (0.792, 0.927, 0.854) | (0.804, 0.987, 0.886) | (0.249, 0.981, 0.397) | (0.469, 0.525, 0.495) |
| EfficientNet | (0.859, 0.987, 0.918) | (0.846, 0.946, 0.893) | (0.870, 0.976, 0.919) | (0.864, 0.933, 0.897) | (0.569, 0.547, 0.558) | (0.575, 0.808, 0.672) |
| ShufflNet | (0.820, 1.000, 0.901) | (0.917, 0.946, 0.931) | (0.816, 0.976, 0.889) | (0.818, 0.960, 0.883) | (0.260, 0.962, 0.410) | (0.561, 0.627, 0.593) |
| MSI-DETECT (best) | (0.831, 0.831, 0.831) | (0.797, 0.887, 0.839) | (0.980, 0.926, 0.952) | (0.692, 0.783, 0.735) | (0.702, 0.755, 0.727) | (0.686, 0.824, 0.749) |
| MSI-DETECT (worst) | (0.674, 0.817, 0.739) | (0.662, 0.924, 0.772) | (0.960, 0.889, 0.923) | (0.630, 0.739, 0.680) | (0.565, 0.660, 0.609) | (0.639, 0.674, 0.656) |
| MSI-DETECT (aggr) | (0.980, 0.769, 0.862) | (0.712, 0.887, 0.790) | (0.962, 0.944, 0.953) | (0.773, 0.739, 0.756) | (0.595, 0.830, 0.693) | (0.669, 0.769, 0.715) |

**Supplementary Table 4**: The MSI prediction performance of the models on the colorectal cancer datasets when the training dataset was Yonsei-1. The numbers in the parentheses in each cell are precision, recall, and F1 score, respectively.

| Model | STMary-GC (precision, recall, F1) | GC-ICI (precision, recall, F1) | Molecular subtypes (precision, recall, F1) |
|---|---|---|---|
| MSI-SEER | (0.500, 0.864, 0.633) | (0.435, 0.714, 0.540) | (0.682, 0.882, 0.769) |
| DGP+CTransPath | (0.536, 0.682, 0.600) | (0.700, 0.500, 0.583) | (0.667, 0.941, 0.780) |
| ResNet | (0.462, 0.818, 0.590) | (0.400, 0.286, 0.333) | (0.556, 0.882, 0.682) |
| EfficientNet | (0.419, 0.818, 0.554) | (0.360, 0.643, 0.462) | (0.909, 0.588, 0.714) |
| ShufflNet | (0.583, 0.636, 0.609) | (0.286, 0.571, 0.381) | (0.538, 0.824, 0.651) |
| MSI (best) | (0.400, 1.000, 0.571) | (0.429, 0.429, 0.429) | (0.867, 0.765, 0.812) |
| -DETECT (worst) | (0.357, 0.909, 0.513) | (0.181, 0.929, 0.302) | (0.727, 0.471, 0.571) |
| (aggr) | (0.368, 0.955, 0.532) | (0.286, 0.429, 0.343) | (1.000, 0.471, 0.640) |

**Supplementary Table 5**: MSI prediction performance on the gastric cancer validation datasets when the training data was the combined data from TCGA-STAD and Yonsei-Classic. The numbers in the parentheses in each cell are precision, recall, and F1 score, respectively.

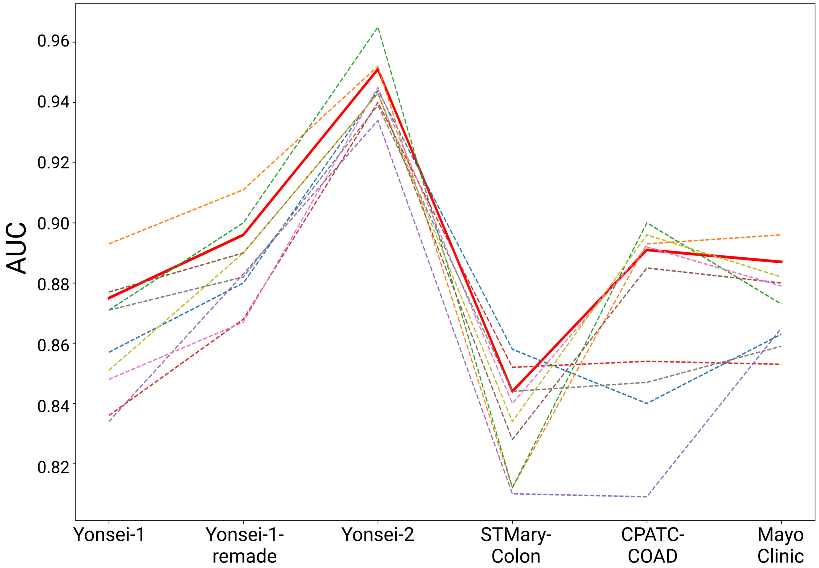


**Supplementary Figure 2**: Prediction performance of each MSIDETECT CNN model for colorectal cancer. Each dashed line represents a separate MSIDETECT CNN model, and the red line represents the aggregated model.

#### A. ROC curves for the methods

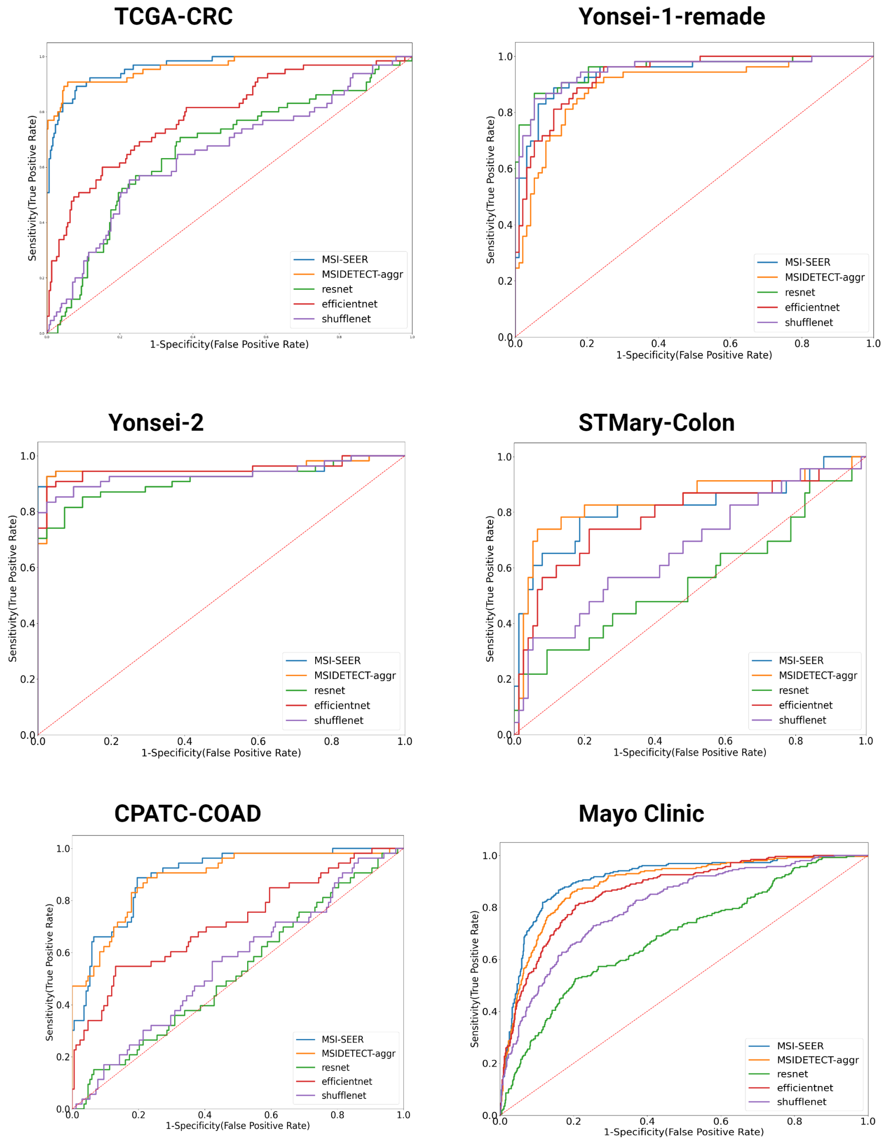


#### B. DeLong's test p-values

| | vs. MSIDETECT | vs. ResNet | vs. EfficientNet | vs. ShuffleNet |
|---|---|---|---|---|
| **TCGA-CRC** | 0.868 | **0.000** | **0.000** | **0.000** |
| **Yonsei-1-remade** | **0.001** | 0.240 | 0.641 | 0.251 |
| **Yonsei-2** | 0.608 | 0.026 | 0.627 | 0.168 |
| **STMary-Colon** | 0.343 | **0.001** | 0.570 | **0.046** |
| **CPATC-COAD** | 0.623 | **0.000** | **0.000** | **0.000** |
| **Mayo Clinic** | **0.001** | **0.000** | **0.000** | **0.000** |

**Supplementary Figure 3**: Comparison of MSI-SEER performance against other models for colorectal cancer. Yonsei-1 was used as the training set. (A) The receiver operating characteristics (ROC) curves of each prediction model for each validation dataset are shown. (B) MSI-SEER was compared to each of the other models with DeLong's test, and the resulting p-values are shown.

**A. ROC curves for the methods**

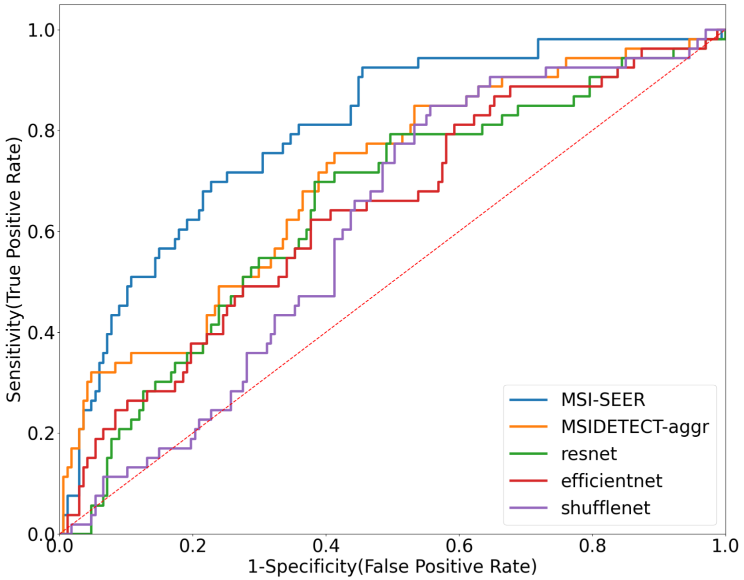


**B. DeLong's test p-values**

| | vs. MSIDETECT | vs. ResNet | vs. EfficientNet | vs. ShuffleNet |
|---|---|---|---|---|
| Combined GC validation data | 0.0066 | 0.0053 | 0.0006 | 0.000 |

**Supplementary Figure 4**: Comparison of MSI-SEER performance against other models for gastric cancer. The combined data (TCGA-STAD and Yonsei-Classic) was used as the training set. For the validation dataset, we chose to combine STMary-GC, Molecular subtypes, and GC-ICI into a single larger dataset because AUC scores can be unreliable when the sample size is small. (A) The receiver operating characteristics (ROC) curves of each prediction model for the validation dataset are shown. (B) MSI-SEER was compared to each of the other models with DeLong's test, and the resulting p-values are shown.

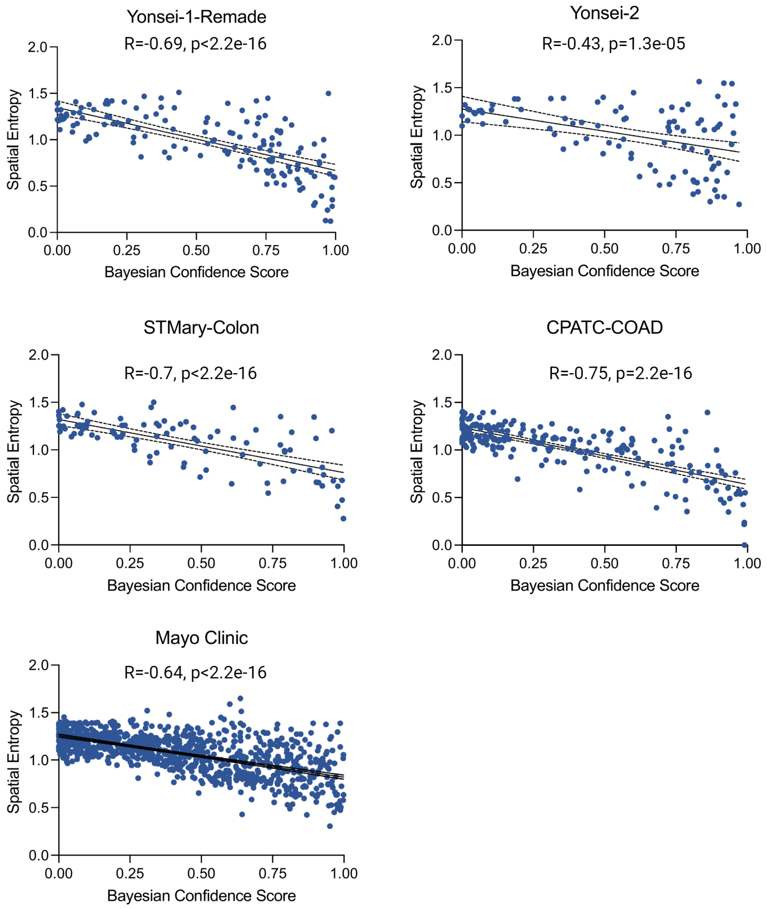


**Supplementary Figure 5**: Correlation between uncertainty and spatial entropy of the test datasets in colorectal cancer. The Pearson correlation coefficients and the corresponding p-values are shown.

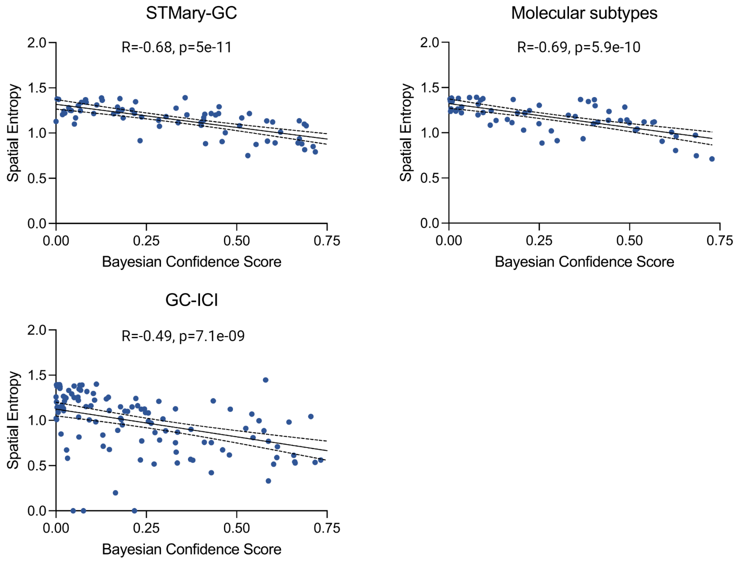


**Supplementary Figure 6**: Correlation between uncertainty and spatial entropy of the test datasets in gastric cancer. The Pearson correlation coefficients and the corresponding p-values are shown.

**Supplementary Table 6**: Prediction improvement in terms of precision, recall, and F1 score in Figure 3 of the Results section: (A) Original Performance and (B) Performance after most uncertain predictions removed.

| | (A) | | | (B) | | |
|---|---|---|---|---|---|---|
| | Precision | Recall | F1 | Precision | Recall | F1 |
| Colorectal | 0.726 | 0.791 | 0.757 | 0.726 | 0.858 | 0.787 |
| Gastric | 0.493 | 0.698 | 0.578 | 0.614 | 0.628 | 0.621 |

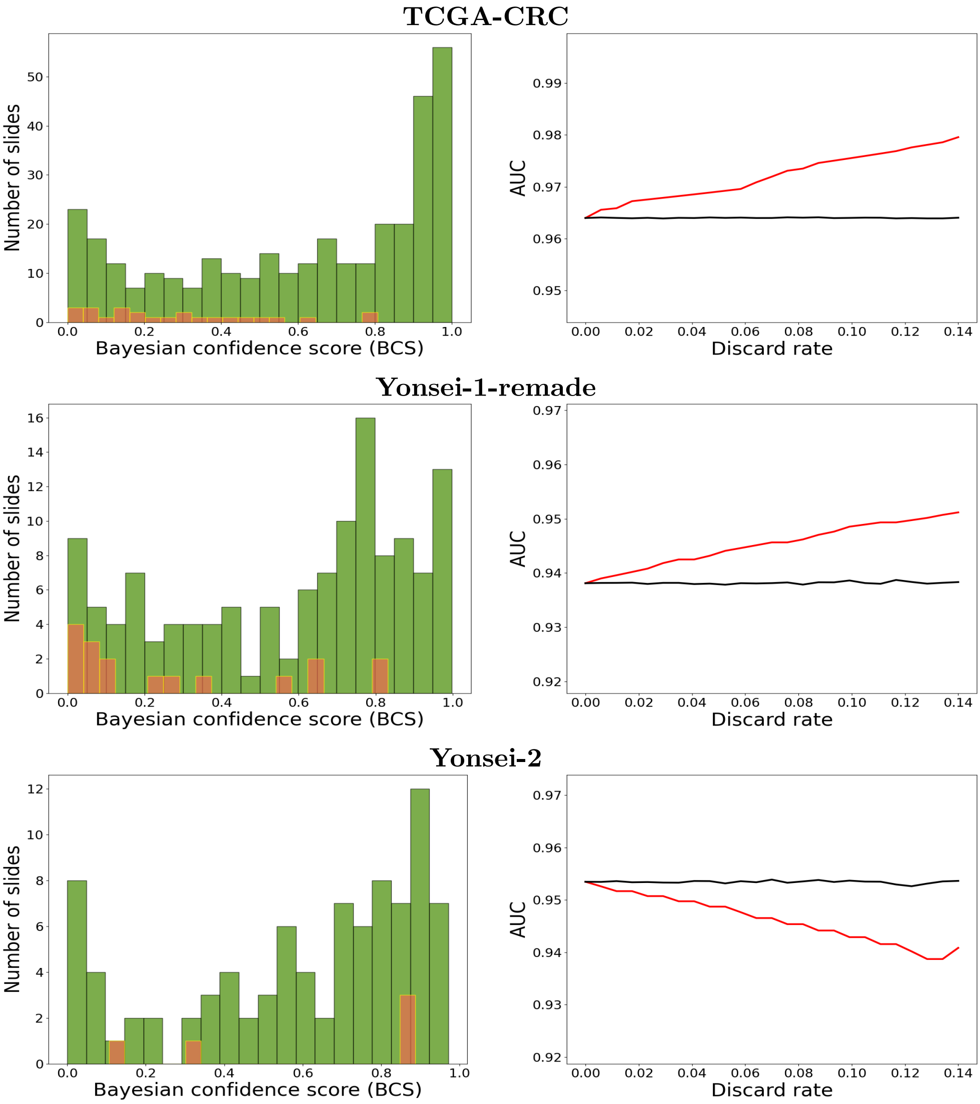
TCGA-CRC
Number of slides
Bayesian confidence score (BCS)
AUC
Discard rate
Yonsei-1-remade
Number of slides
Bayesian confidence score (BCS)
AUC
Discard rate
Yonsei-2
Number of slides
Bayesian confidence score (BCS)
AUC
Discard rate

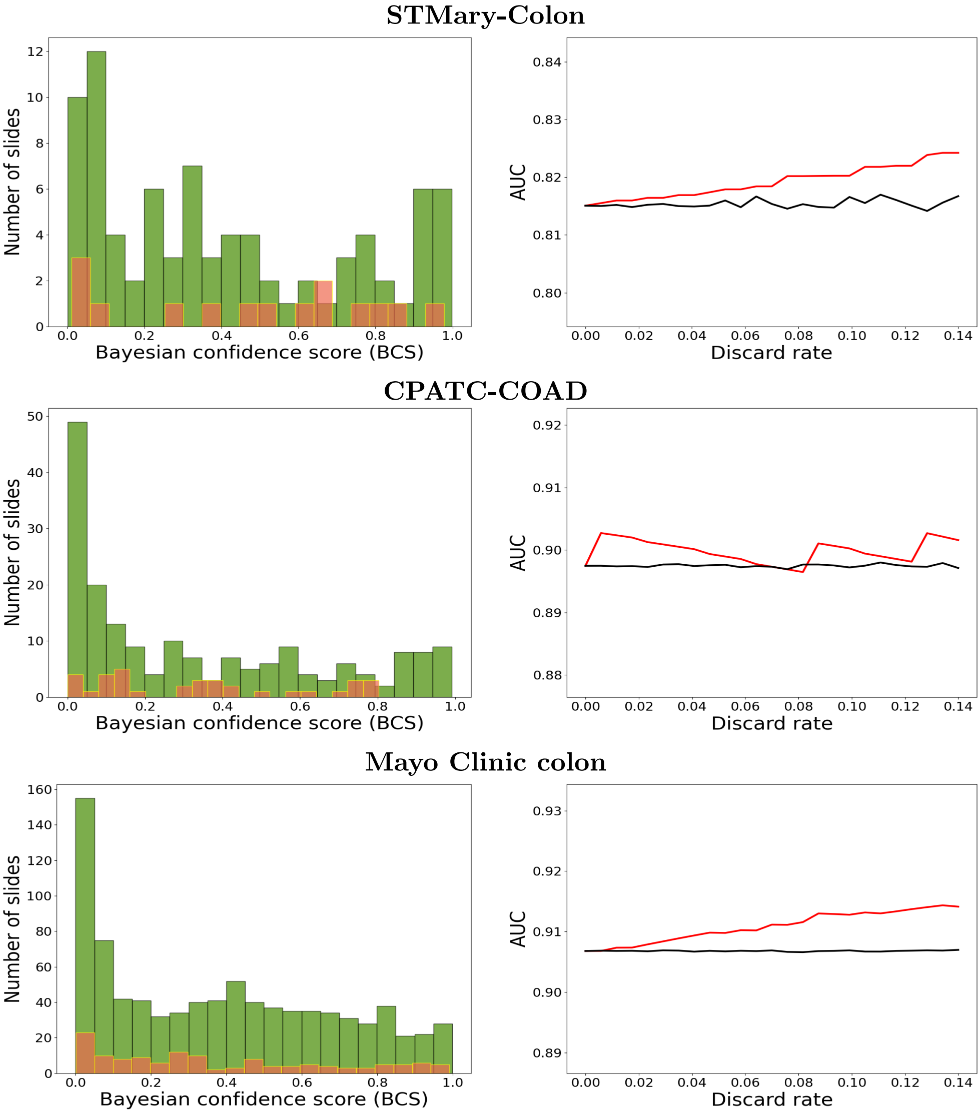


**Supplementary Figure 7**: Excluding the most uncertain predictions improves WSI prediction performance in colorectal cancer. For the histograms, green represents correctly classified WSI and orange represents incorrectly classified WSI. The red line represents the change in the performance when the most uncertain predictions (as measured by the BCSs obtained by our model) were discarded while the black line is the average change in the performance the predictions are randomly discarded 1,000 times at each rate. Discard rate: discarded WSIs/total WSIs.

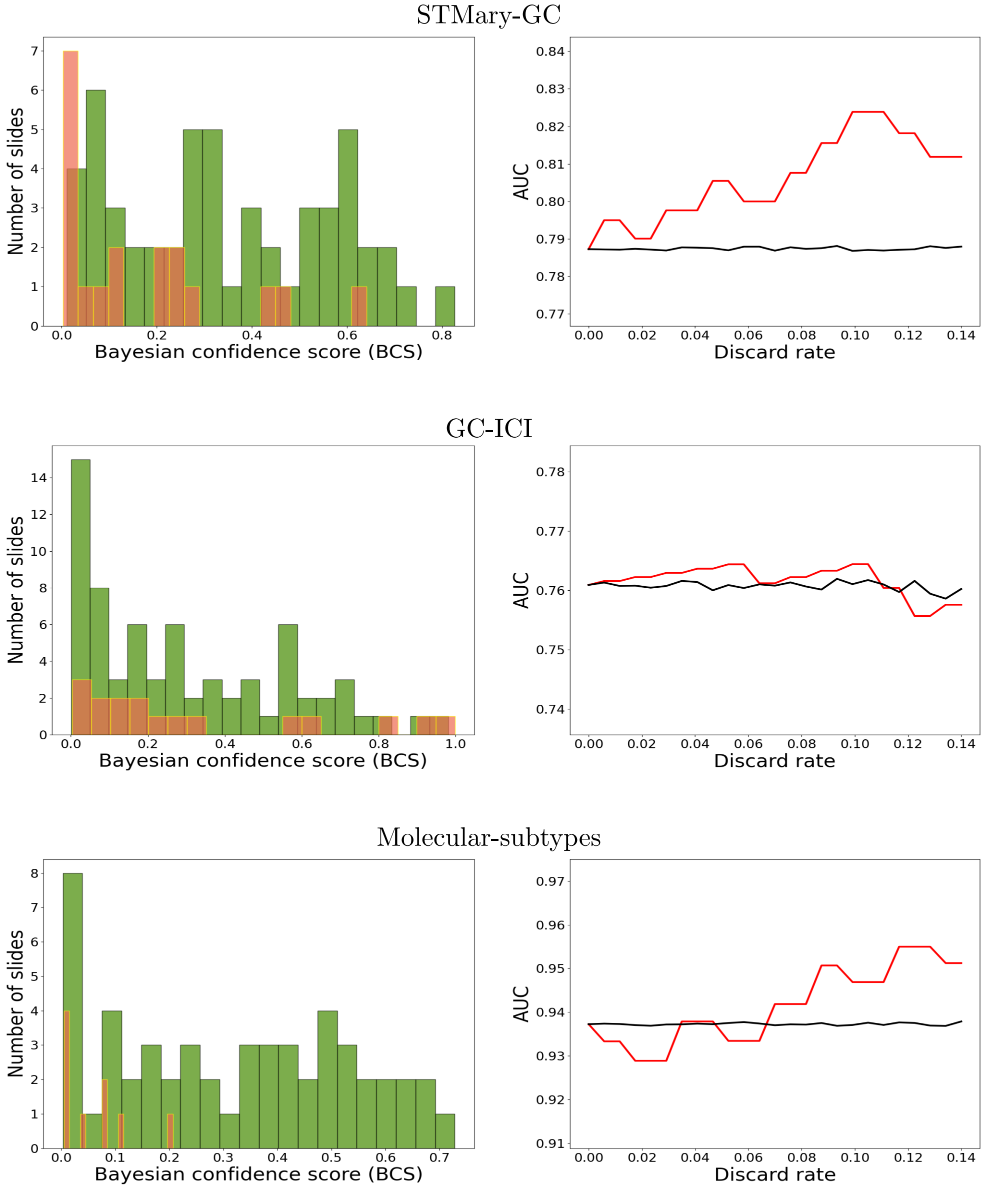


**Supplementary Figure 8**: Excluding the most uncertain predictions improves WSI prediction performance in gastric cancer. For the histograms, green represents correctly classified WSI and orange represents incorrectly classified WSI. The red line represents the change in the performance when the most uncertain predictions (as measured by the BCSs obtained by our model) were discarded while the black line is the average change in the performance the predictions are randomly discarded 1,000 times at each rate. Discard rate: discarded WSIs/total WSIs.

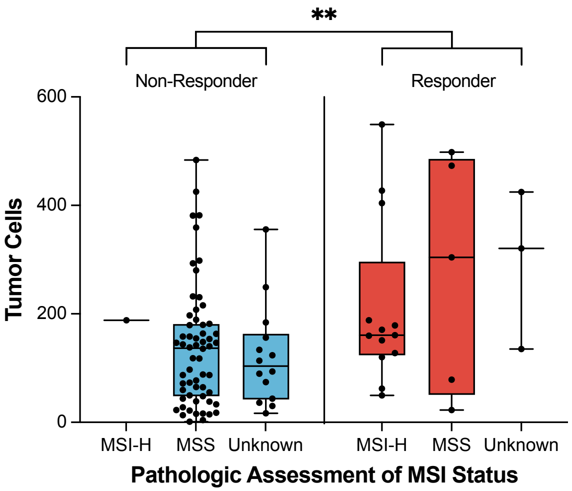


**Supplementary Figure 9**: Comparison of tumor cell counts in the predicted MSI-H tumor region in gastric cancer patients treated with immune checkpoint inhibitors, stratified by treatment response. All responders were compared to all non-responders. $**\ P < 0.01$, by Wilcoxon test.

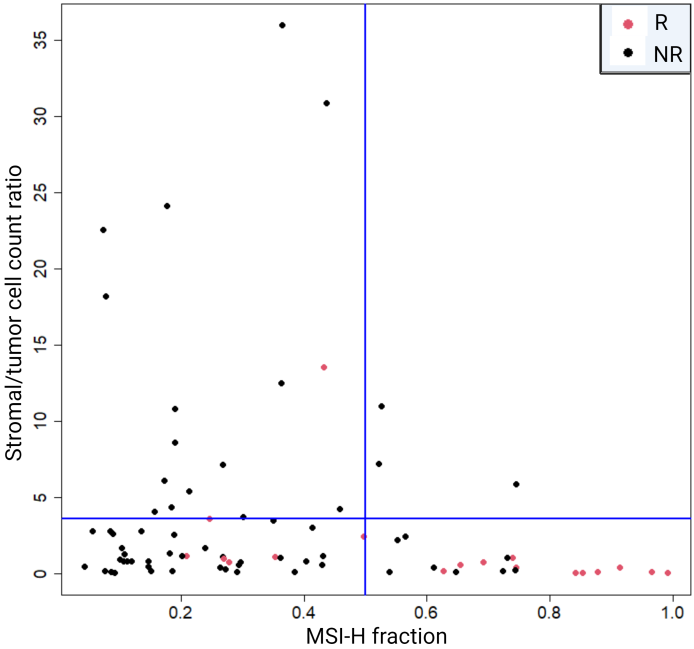


**Supplementary Figure 10**: Scatter plot of MSI-H fraction versus stroma-to-tumor ratio for samples with MSI status information (as training data). Each point represents one slide. The vertical line is at 0.5 (the cutoff for the MSI-H fraction). This value was determined as the cutoff for standard binary classification problems. The horizontal line is at 3.6 (the cutoff for the stroma-to-tumor ratio). This value was determined as the largest value of the stroma-to-tumor ratio in the responder group after excluding the outlier.

**Prediction Based on MSI Status and MSI-SEER Predictions for Samples with Unknown MSI Status**

**Actual**

| **Predicted** | Responder | Non-Responder | |
|---|---|---|---|
| Responder | 2 (True positive) | 0 (False positive) | Positive Predictive Value 100% |
| Non-Responder | 1 (False negative) | 14 (True negative) | Negative Predictive Value 93.3% |
| | Sensitivity 66.7% | Specificity 100% | |

**Supplementary Figure 11**: Predicting gastric cancer response to immune checkpoint inhibitors. The prediction rules were derived based on predicted MSI-H fraction by MSI-SEER and the stroma-to-tumor ratio in the predicted MSI-H areas of samples with MSI status information. Samples with unknown MSI status were used as test data. The confusion matrix with the classification performance evaluation metrics is presented.
